# Network Neuroscience of Human Multitasking: Local Features Matter

**DOI:** 10.1101/2025.04.30.651494

**Authors:** Marie Mueckstein, Kirsten Hilger, Stephan Heinzel, Urs Granacher, Michael A. Rapp, Christine Stelzel

**Affiliations:** International Psychoanalytic University Berlin Germany; Universität Potsdam Germany; Universität Würzburg Germany; Freie Universität Berlin Germany; TU Dortmund University Dortmund Germany; University of Freiburg Freiburg i.B. Germany

**Keywords:** Functional Connectivity, Dual task, Crosstalk, Executive Control, Modality Compatibility

## Abstract

The neural basis of multitasking costs is subject to continuing debate. Cognitive theories assume that overlap of task representations may lead to between-task crosstalk in concurrent task processing and thus requires cognitive control. Recent research suggests that modality-based crosstalk contributes to multitasking costs, involving central overlap of modality-specific representations. Consistently increased costs for specific modality pairings (visual-vocal and auditory-manual vs. visual-manual and auditory-vocal) were demonstrated (modality-compatibility effect), which were recently linked to representational overlap in the auditory cortex. However, it remains unclear whether modality-based crosstalk emerges from overlapping patterns of global brain connectivity and whether resolving it requires additional involvement of cognitive control as reflected in the fronto-parietal control network. This preregistered functional imaging study investigates these questions in 64 healthy, young human adults. Specifically, we focus on the modality-compatibility effect in multitasking by employing functional connectivity (FC) analysis. First, we tested the FC similarity between the single-task networks. Second, we compared the strength of the control network in whole-brain FC between dual tasks. We found no evidence for different FC similarities of single-task networks between modality pairings and no additional involvement of the control network during dual tasks. However, post-hoc connectivity analysis revealed a brain-behavior correlation for the modality-compatibility effect in dual tasks. This effect was locally restricted to FC between lateral frontal and sensory auditory regions, providing evidence for the modality-based crosstalk theory. More generally, the findings suggest that robust behavioral differences in multitasking are not necessarily related to global functional connectivity differences but to local connectivity changes.

**Significance:** Our lives are dominated by multitasking. Understanding the neurocognitive mechanisms of multitasking and its associated limitations is relevant for safety-relevant consequences. Here, we investigate functional brain-connectivity patterns associated with modality-based crosstalk in multitasking, describing unintentional exchange of information between two tasks, leading to robust dual-task costs. Our network analyses revealed no significant difference between single or dual tasks with varying degrees of modality overlap on the whole-brain level. However, specific connections between cognitive control-related frontal and sensory-related regions were associated with individual multitasking performance, emphasizing that behavioral differences can arise from specific neural interactions rather than from widespread network reconfigurations. These findings advance our understanding of the underlying mechanisms responsible for modality-based crosstalk, providing practical implications for human-machine interactions.

## Introduction

Whether it is cooking while listening to a podcast or driving while talking on the phone - multitasking is omnipresent in everyday life, even though it usually comes with performance costs. Neuroimaging studies investigating the neural basis of performing two tasks concurrently (i.e., dual-tasking) show consistent activity in frontoparietal regions associated with cognitive control (meta-analysis by Worringer et al., 2019). This activity occurs when comparing dual-task to single-task activity and dual tasks with different cognitive demands. Cognitive control may also be involved in processing representational overlap between tasks in a dual-task context. Recently, several functional imaging (fMRI) studies (Alavash et al., 2015; Mueckstein et al., 2025; Nijboer et al., 2014; Paas Oliveros et al., 2023) complemented the available behavioral research on the role of representational overlap for dual-task costs (Janczyk et al., 2014; Koch, 2009). A widespread assumption is that representational overlap causes central crosstalk, i.e., the unintended exchange of information between tasks, thus negatively affecting performance (Lien & Proctor, 2002; Logan & Gordon, 2001; Navon & Miller, 1987).

Recently, Mueckstein et al. (2025) investigated the neural basis of modality-based crosstalk - a specific type of crosstalk, where central overlap between modality-specific information is assumed to cause dual-task costs. Several behavioral studies provided evidence for robustly increased dual-task costs when pairing visual-vocal (VV) with auditory-manual (AM) tasks (i.e., modality-incompatible) compared to visual-manual (VM) with auditory-vocal (AV) tasks (i.e., modality-compatible) (Göthe et al., 2016; Hazeltine et al., 2006; Mueckstein et al., 2022; Stelzel et al., 2006). According to the modality-based crosstalk account, this modality-compatibility effect in dual tasks arises from the overlap between the stimulus modality in one task (e.g., auditory stimulus) and the anticipated sensory action consequence (e.g., auditory action effect of a vocal response) in the simultaneously performed tasks (Figure 1B) (Schacherer & Hazeltine, 2020, 2021, 2023).

**Figure 1.**
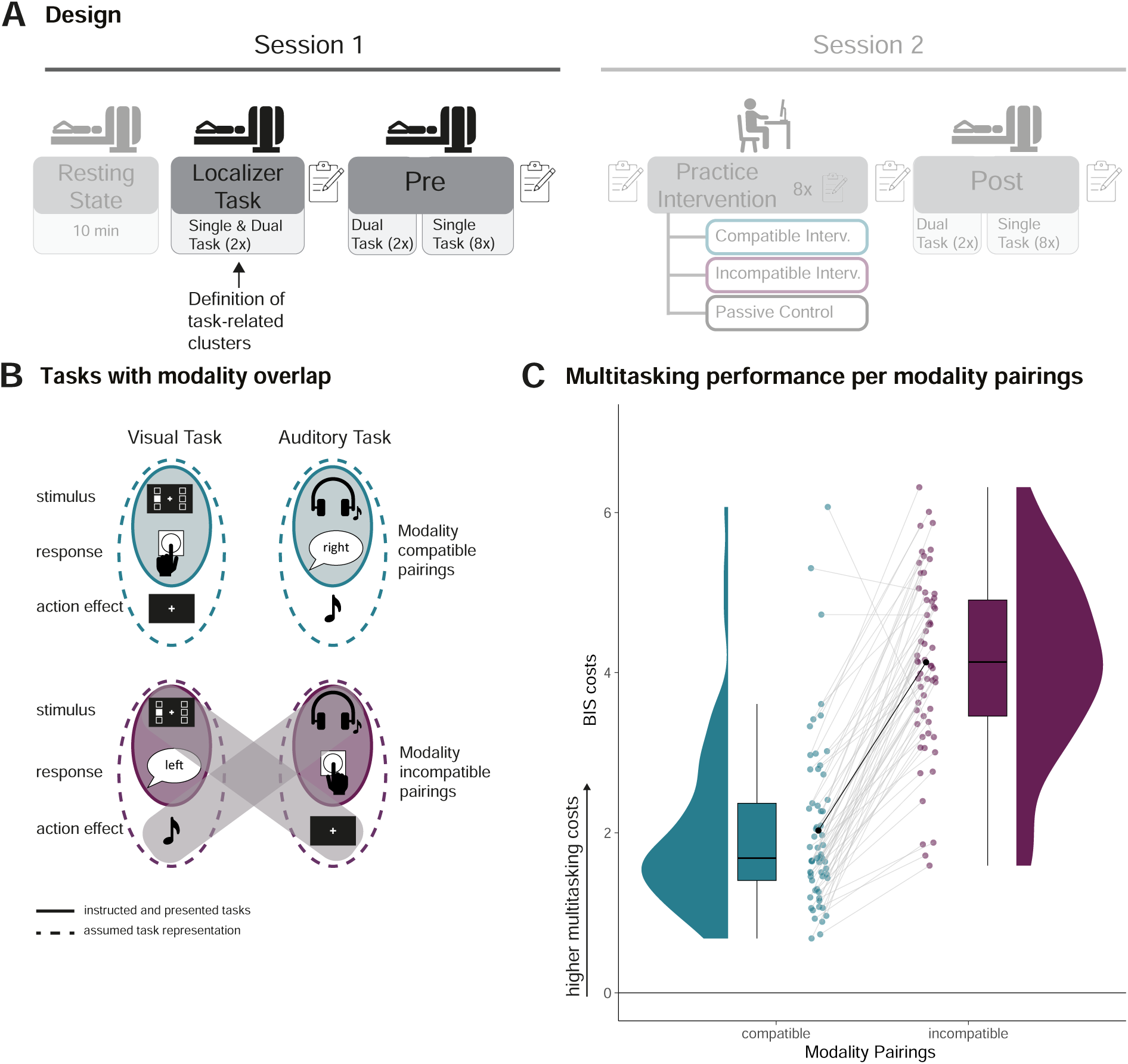
Experimental design, tasks and behavioral results. **A**: Overview of the two fmri sessions. The parts of the study that are not relevant here are hidden and will be reported elsewhere. Localizer tasks are the same tasks as used in the Pre part in both modality pairings and for single and dual task in a block-design (two runs). This data is used to define the univariate task-activity clusters. The main part (Pre) consists of two dual-task runs, one per modality pairing. Followed by eigth single task runs, each containing eigth blocks one for each single task type (task complexity x modality pairing). **B**: The upper part of the figure depicts the stimulus-response pairings for the modality-compatible pairing, comprising a visual stimulus with a manual response and an auditory stimulus combined with a vocal response. For each response, the corresponding natural action effects are depicted as well. Note that the action effect of the manual response is not exclusively visual but also somatosensory. Likewise, the action effect of the vocal response is typically auditory (i.e., hearing oneself speaking) but also somatosensory (i.e., feeling one’s mouth move). The lower part depicts the modality-incompatible pairing. The visual stimulus is paired with a vocal response and the auditory stimulus with a manual response. In this condition, the match between action-effect modality and stimulus modality is between tasks, potentially causing interference when presented as dual task, due to higher overlap. **C**: The graph provides the distribution, boxplot, mean, and individual performance per modality pairing of the balanced integrated score (BIS) cost parameter. Multitasking cost is the difference between dual task and single tasks, measured as BIS, which is an integration of reaction times and accuracies (BIS = 0 means no difference between single and dual task performance). The graph demonstrates the robustly higher multitasking cost for the modality-incompatible pairing, compared to the modality-compatible pairing.

Mueckstein et al. (2025) provided neural evidence for this hypothesis by applying multi-voxel pattern analysis to fMRI data. Between-task overlap in modality-incompatible single tasks was exclusively present in the auditory cortex, supporting the role of local modality-specific representational overlap for dual-task performance. Additionally, a region-of-interest (ROI) analysis on univariate fMRI data revealed greater involvement of the left inferior frontal sulcus during the performance of modality-incompatible compared to modality-compatible dual tasks, suggesting higher cognitive control demands for modality-incompatible pairings (Stelzel et al., 2006).

However, the human brain is more than a collection of circumscribed regions operating in isolation. Network neuroscience highlights the importance of dynamic interactions (Rubinov & Sporns, 2010) and stresses the importance of considering whole-brain connectivity (Sporns, 2011). Yet, little is known about these interactions in dual-task processing, including the processing of modality-based crosstalk. Our study closes this gap. First, we compared whole-brain single-task functional connectivity (FC) patterns between modality pairings. Alavash et al. (2015) demonstrated that greater overlap between single-task modules predicts higher dual-task costs. Combining this finding with the results of Mueckstein et al. (2025), we hypothesized that the single-task FC pattern overlaps more strongly for modality-incompatible pairings than for modality-compatible pairings. Second, we compared dual-task-related FC patterns between modality pairings to investigate whether the connectivity of fronto-parietal regions associated with cognitive control processes in dual tasks (Worringer et al., 2019) differ.

Based on the previous univariate data (Stelzel et al., 2006), we hypothesized larger involvement of the control network (Yeo et al., 2011) in modality-incompatible dual tasks related to resolving modality-based crosstalk.

We observed no significant differences in FC between modality pairings, neither between the single-task similarity of FC patterns nor in the involvement of the control network between dual tasks. However, post-hoc analyses revealed higher local FC between lateral frontal regions and auditory regions in the modality-incompatible compared to the modality-compatible dual task. This difference in FC was associated with individual differences in dual-task performance. Our results thus provide evidence for the assumption that controlling the representational overlap of modality-specific information by engaging lateral frontal brain regions is relevant for successfully resolving modality-based crosstalk in multitasking.

## Methods

### Preregistration

This study is part of a larger research project (see Figure 1A). The project was pre-registered before any data analysis took place (https://osf.io/whpz8) and contains two different methodological approaches using the same sample. The first was a multi-voxel pattern analysis (MVPA) reported in Mueckstein et al. (2025), and the second is reported here. Accordingly, sections in the methods are mostly copied from the preregistration and shortened. Note that any deviations from the preregistration are explicitly marked as such.

### Participants

In total, 71 healthy, right-handed adults aged 18 to 30 with German as their first language (or comparable level) and normal or corrected-to-normal vision participated in the study. We recruited participants via flyers and mailing lists of local universities and online advertisements. Exclusion criteria were any neurological or psychiatric diseases, current medical conditions that could potentially influence brain functions, past or present substance abuse (alcohol and drugs), a self-reported weakness in distinguishing left and right, and common contraindications for MRI scanning. Table 1 provides the final sample size along with age and gender distributions per analysis. We provide a detailed overview of why participants were excluded and how many participants were part of both analyses (MVPA and connectivity) in the supplementary material (Table S2). The specific criteria for exclusion are reported in the sections “Behavioral Tasks” and “Function Brain Connectivity”. All participants provided written informed consent before the study started and were reimbursed with 60 € or course credit. The ethics committee of the Freie Universität Berlin approved the study (approval number: 018.2021) following the latest version of the Declaration of Helsinki.

**Table 1.**
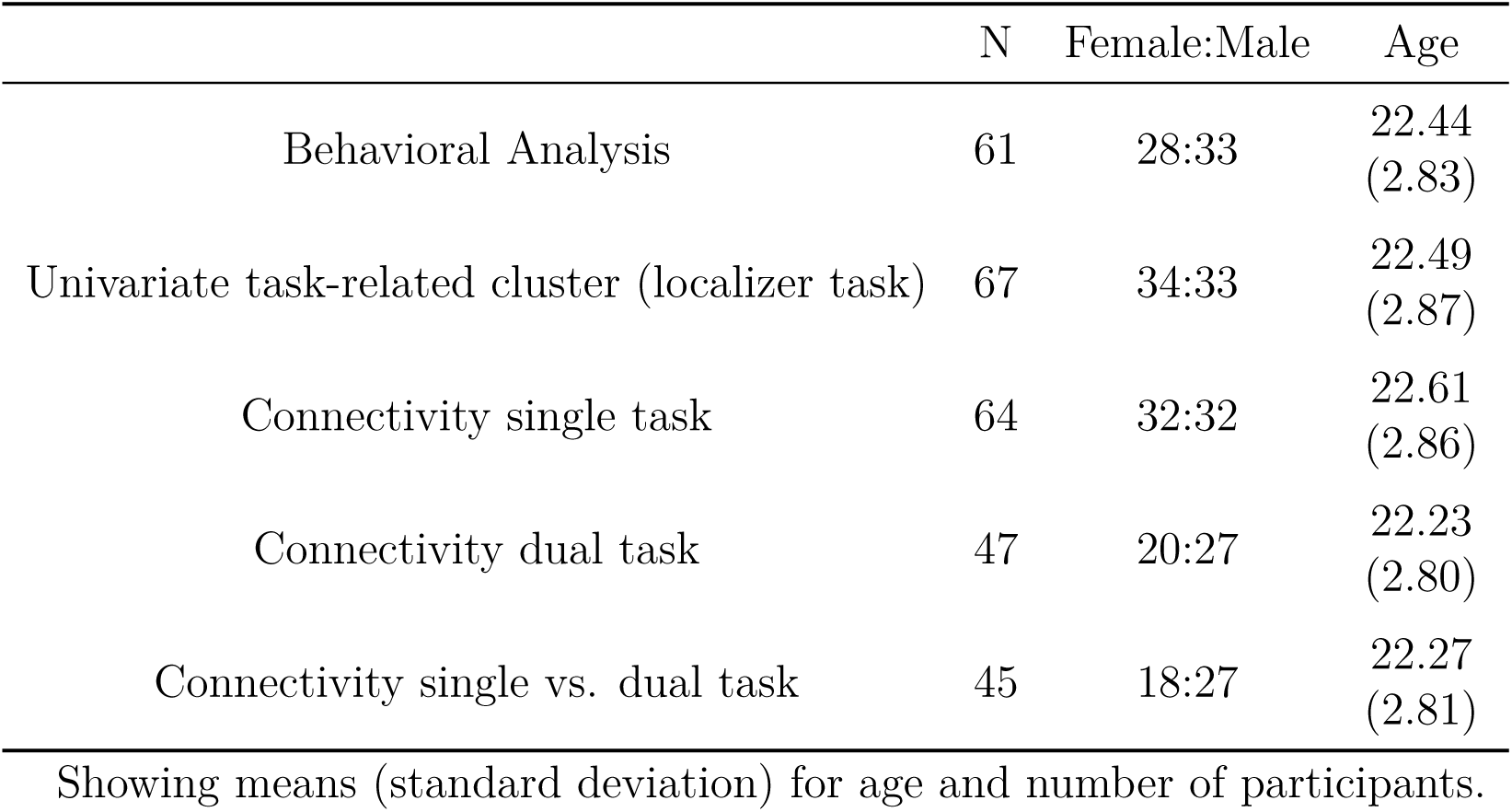
Participants age and sex ratio per analysis.

|  | N | Female:Male | Age |
| --- | --- | --- | --- |
| Behavioral Analysis | 61 | 28:33 | 22.44<br>(2.83) |
| Univariate task-related cluster (localizer task) | 67 | 34:33 | 22.49<br>(2.87) |
| Connectivity single task | 64 | 32:32 | 22.61<br>(2.86) |
| Connectivity dual task | 47 | 20:27 | 22.23<br>(2.80) |
| Connectivity single vs. dual task | 45 | 18:27 | 22.27<br>(2.81) |

Showing means (standard deviation) for age and number of participants.

### Experimental procedure

Participants completed one online session and two fMRI sessions in the context of a practice intervention design. Here, we will focus on the results of fMRI session one (Figure 1A). Due to high drop-out rates for the second (post-practice) session and the strict head movement criteria for connectivity analyses (see below), the sample size was not large enough to analyze the practice-related changes between the two sessions with sufficient power as it was initially planned in the pre-registration (39 participants left, with 10, 13 and 16 per intervention group for the dual task).

The online session consisted of behavioral and cognitive measures and is part of a different study with a separate pre-registration (https://osf.io/nfpqv). The two fMRI sessions (each 2.5 -3 h) took place at the Cognitive Center for Neuroscience Berlin (CCNB). During the first session, participants started outside the scanner with a short familiarization of the modality-pairing tasks (256 trials, 32 per single task, 64 per dual task). The following in-scanner part started with a ten-minute resting-state measurement with open eyes. It was followed by two runs of a localizer task, each containing single and dual tasks in both modality pairings, organized in a block design. Each run of the localizer task comprised six blocks, each containing 16 trials. The following two runs included only dual-task trials. Each run consisted of 128 trials and was assigned to one modality pairing (i.e., modality-compatible or modality-incompatible). The remaining eight runs contained only single-task trials in both modality pairings, with an easy and a difficult version of the task. The manipulation of task difficulty served as a control analysis for the MVPA and is not distinguished here. In each run, every task, modality pairing, and difficulty combination occurred only once, resulting in eight blocks with 16 trials per block. A visual overview of the run and task structure is presented in the supplementary material ( Figure S1). Details about the second session can be found in the pre-registration of the study (https://osf.io/whpz8) and in Mueckstein et al. (2025).

### Behavioral tasks

Participants completed sensorimotor choice reaction tasks, as single and dual tasks with two different modality pairings (modality-compatible and modality-incompatible, compare Figure 1B). The visual stimuli were a white square (pixel size 56.8 x 56.8) on a black background next to a fixation cross (pixel size 41.1 x 41.1, thickness 9.9 pixel) at six different positions (top, center, bottom, either right or left from the fixation cross). The auditory stimuli were pure tones at different frequencies (200, 450, and 900 Hz), presented on either the right or left ear. In the single-task blocks, only one stimulus per trial was presented, while in the dual-task blocks, one visual and one auditory stimulus were presented simultaneously (stimulus-onset-asynchrony = 0 ms). All stimuli were presented for 200 ms, followed by a response interval of 1500 ms and an inter-stimulus interval of 200 ms. Each run concluded with an 8 s fixation period.

Participants were asked to indicate the side on which the stimulus was presented by either pressing a button with their right or left hand (index finger) and/or by saying the German word for “right” or “left”. The pairing of visual-manual and auditory-vocal is regarded as modality-compatible, and the pairing of visual-vocal and auditory-manual as modality-incompatible. Consequently, within each dual-task condition, none of the pairings had a direct overlap between the presented stimulus modality and the required response modality. Nevertheless, they overlap between stimulus modality and the modality of the anticipated response-related sensory consequences in the modality-incompatible condition.

During the single-task runs, we manipulated the task difficulty by adding visual noise, increasing the distance of the stimulus from the center, and reducing the contrast between the stimulus and background. Similarly, we added white noise for the auditory stimulus and reduced the volume. As preregistered, only easy blocks were used for the behavioral analysis to better compare the results to earlier studies. We did not distinguish between easy and difficult single-task blocks for the fMRI analysis.

The order of the dual-task runs was counterbalanced across participants, while the block position during each single-task run was randomized for each participant. Each stimulus was presented equally often and in random order per trial within each block.

Data exclusion was determined per run. Specifically, participants were excluded if one of the following criteria applied to either one dual-task run, more than one localizer run, or more than three single-task runs: On a behavioral level, data were excluded if participants used the wrong response modality in a block for more than five trials (e.g., vocal response in VM block) or committed more than 30 % errors (including omissions) in the single-task or localizer runs. Due to the high error rate in the dual-task runs (whole sample (N=71) modality-compatible, *M* = 19.68% [*SD* = 16.11, max = 73.44 %], modality-incompatible *M* = 42.48 % [*SD* = 17.28, max = 73.44 %]) we deviated from the pre-registered protocol and applied the 30% criteria only to trials in which both stimuli required the same response location (i.e., congruent trials, averaged for both modality pairings in a dual-task trial). This ensured that participants generally understood the task.

Further, we decided to deviate from the pre-registration in the analysis of behavior. Specifically, we used a balanced integrated score (BIS, Liesefeld & Janczyk, 2019; Mueckstein et al., 2025), which is defined as the difference between the z-standardized reaction times and accuracies instead of separately analyzing reaction times and error rates. Liesefeld and Janczyk (2019) showed that the BIS parameter controls well for speed-accuracy trade-offs, thus accounting for different individual response strategies (i.e., focusing either more on accuracy or speed). Additionally, it reduces the number of analyses, which increases the statistical power and clarity of analyses. We used a standard paired t-test together with a Bayesian paired t-test to analyze the behavioral data.

Statistical analysis and plotting were conducted in R (version 4.2.2, R. C. Team, 2018) with RStudio (version 2023.12.1, Rs. Team, 2019), the tidyverse package (version 2.0.0, Wickham et al., 2019) and the BayesFactor package (version 0.9.12, Morey & Rouder, 2024). Brain-related plots were created with the ggseg package (version 1.6.5, Mowinckel & Vidal-Piñeiro, 2019), the brainconn package (version 0.1, Chopra et al., 2023) and one function from the CONN Toolbox (version 22a, Nieto-Castanon & Whitfield-Gabrieli, 2022). The manuscript was created with the papaja package (version 0.1.2, Aust & Barth, 2023).

### MRI data acquisition & preprocessing

#### Acquisition

Due to a scanner upgrade at the imaging center, the data were acquired with two different MRI scanners. The first 25 participants were measured with Siemens Magnetom TIM TRIO syngo 3T and the remaining participants with Siemens Magnetom 3.0T Prisma both with a 32-channel head coil and based on the same parameters. At the end of the session, a high-resolution T1-weighted structural image was measured with 176 interleaved slices, 1 mm isotropic voxels; TE = 2.52 ms, TR = 1900 ms, FoV = 256 x 256 x 176 mm. The functional runs consisted of 139 whole-brain echo-planar images of 37 interleaved slices for the localizer task, and the dual-task runs and 183 whole-brain echo-planar images for each single-task run. Each functional run was acquired with 3 mm isotropic voxels, TE = 30 ms, TR = 2000 ms, flip angle =75°, FoV = 192 x 192 x 133 mm. After each dual-task run, a grey-field mapping was measured (3 mm isotropic voxel, TE1 = 4.92 ms and TE2 = 7.38 ms; TR = 400 ms; FoV = 192 x 192 x 133 mm, flip angle = 60°). Participants received auditory stimuli via MRI-compatible headphones (Sensi-Metrics S14, SensiMetrics, USA). Visual stimuli were projected on a screen at the end of the bore, which participants could view through a mirror attached to the head coil. Vocal responses were recorded via an MRI-compatible microphone (OptimicTM MEG, Optoacoustics, Israel) and manual responses via MRI-compatible 4-button bimanual boxes (HHSC-2x2, Current Designs, USA).

#### Pre-processing

We converted the fMRI DICOM data into BIDS format, using dcm2bids (version 2.1.6, Boré et al., 2023) and applied the preprocessing pipeline of fMRIprep (version 21.0.2, Esteban et al., 2019), comprising of 3D motion correction and slice-time correction.

Pre-processing contained the alignment of all functional data to a generated reference image, co-registration, and the transformation to standard space. Anatomical T1 images were transformed into standard MNI space. Please find a detailed description of the pre-processing in the files generated by fMRIprep in the preregistration (https://osf.io/whpz8).

### fMRI data analysis

#### Univariate region-of-interest definition

We conducted the first-level analysis in SPM12 on the two normalized and smoothed (8 mm FWHM Gaussian) BOLD runs of the localizer tasks with a block design to define activity-based task-related clusters. We included six motion parameters (3x rotation, 3x translation) and the framewise displacement parameter as regressors of no interest in the general linear model. We calculated the statistical parametric maps for each participant, including contrasting the stimulus modalities (visual vs. auditory), the response modalities (manual vs. vocal), and the task type (single vs. dual task). On the second level, individual brain maps were averaged and tested voxel-wise using a one-sample t-test between the defined contrasts. We used a cluster-wise FWE-corrected significant threshold of *p* = .05 on the voxel level. For the single vs. dual-task contrast, we restricted the cluster selection to the frontal lobe, as previous studies on dual-tasking consistently showed frontal activity using the same contrast. This process resulted in one task-based activity cluster per hemisphere for visual, auditory, manual, vocal, and dual tasks (frontal). In case there was more than one remaining cluster, we selected the one with the maximum intensity.

#### Preprocessing functional brain network connectivity

We followed the preprocessing steps described by Thiele et al. (2022), and in that the recommendations of Parkes et al. (2018) (pipeline No.6). The final bold time series were created by using the “app-fmri-2-mat” code locally, provided by Josh Faskowitz (https://github.com/faskowit/app-fmri-2-mat). In this code, the nuisance regression strategy using the parameters calculated by the fMRIprep confound file was applied. For the following parameters, the raw values along with all three derivatives were used: global signal, cerebrospinal fluid, white matter, 3x translation, 3x rotation, and framewise displacement. In addition, mean task-evoked activation was removed using the basis-set task regressors as suggested by Cole et al. (2019). A task run was only included in the analysis if the mean framewise displacement was below 0.2 mm, the proportion of spikes larger than 0.25 mm was below 20 %, and if there were no spikes above 5 mm (Parkes et al., 2018; Thiele et al., 2022). In addition, the behavioral performance criteria described above (“Behavioral tasks”) must also be fulfilled.

Deviating from the pre-registration, we applied the schaefer200 instead of the schaefer100 parcellation (Schaefer et al., 2018) for more fine-grained insights. Note that both parcellations showed the same results for the two pre-registered analyses. The results with the pre-registered parcellation schaefer100 can be found in the supplementary material (Table S22 - S27). All schaefer200 regions were assigned to one out of seven functional brain networks (Yeo et al., 2011): visual, somatomotor, dorsal attention, saliency-ventral attention, limbic, control, default.

The resulting time series per run and participant for the single tasks were then filtered for each single task and averaged over runs to account for potential differences between runs. Functional connectivity matrices (FC) were computed as Fisher z-transformed Pearson correlations between all 200 regions. Accordingly, the Pearson correlation defines the strength of the connection between two regions. This procedure resulted in six different FC matrices: visual-manual, auditory-vocal, visual-vocal, auditory-manual, modality-compatible dual task, and modality-incompatible dual task. Figure 2 provides an overview of the preprocessing steps.

**Figure 2.**
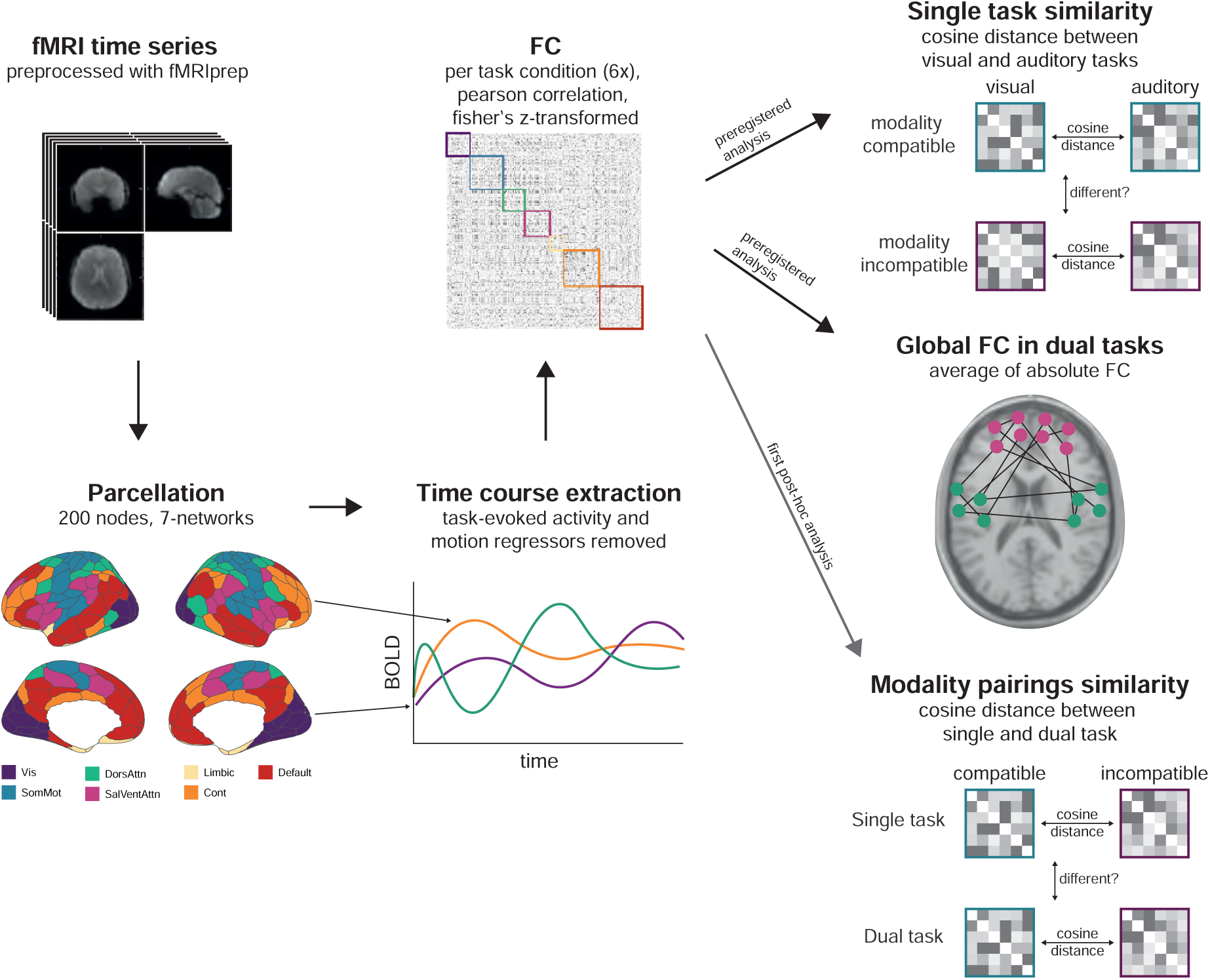
Processing pipeline from preprocessed data to functional connectivity matrix with the two preregistered analyses and one of two post-hoc analysis.

#### Similarity of whole-brain functional connectivity

To investigate the first research question whether whole brain single-task connectivity patterns differ between modality pairings, we employed the cosine distance as a measure of the similarity between FCs. This measure represents the angle (cosine) between two vectors without considering a potential difference in length (Han et al., 2012): 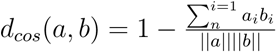, where *a* and *b* are connection vectors and *n* is the total number of connections (compare Thiele et al., 2022).

We compared the brain FC similarity between the two modality-compatible single tasks (visual-manual and auditory-vocal) with the brain FC similarity of the two modality-incompatible single tasks (visual-vocal and auditory-manual). Participants were excluded from this analysis if they showed extensive head movement in more than six (out of eight) single-task runs.

#### Global function connectivity analysis

For each of the dual-task matrices, we calculated the whole-brain FC as the average of the absolute connections within and between all network combinations, while we averaged the between connections for each network (resulting in seven within and seven between network combinations). For each of the network combinations, we applied a linear mixed-effects model, fitted with log-likelihood maximization, with participant and modality pairing as random effects (nlme package, function *lme*, version 3.1-160, Pinheiro et al., 2022). This allows us to assess the difference between modality pairings, while controlling for the factors age, gender and head movement. Additionally, we calculated the Bayesian Factor, using a paired t-test between the two modality pairings, for the seven within and the 21 between-network combinations.

#### Post-hoc functional similarity analysis

Post-hoc (not pre-registered), we averaged the single-task matrices for each modality pairing to further compare the FC similarity between dual-task modality pairings with the FC similarity of the single-task modality pairings (see Figure 2).

For both post-hoc and pre-registered FC similarity analysis, we calculated the FC similarity for each within and between functional network pairings. The between-network combinations were averaged per network, resulting in seven within-network and seven between-network combinations. The same model and additional analysis as for the dual-task matrices were calculated for the FC similarity, by controlling for the number of valid runs in the single tasks.

#### Post-hoc raw functional connectivity analysis and correlation

As additional post-hoc analyses, we calculated the difference between the FC matrix of the modality-incompatible dual task and the modality-compatible dual task per participant. The resulting 200x200 difference matrix was tested against zero, applying a Bayesian one-sample t-test (*ttestBF* function in the r-package BayesFactor) and filtered for connections with a Bayes Factor above 3, which indicates moderate evidence for the alternative hypothesis (Andraszewicz et al., 2015). See Figure 3 for a visual overview of these steps. As we were mainly interested in task-specific differences, only connections between regions that overlapped with the univariate task-relevant cluster from the localizer task were considered. The supplementary material reports the whole brain FC matrix filtered only by the Bayes Factor (compare Figure S2).

**Figure 3.**
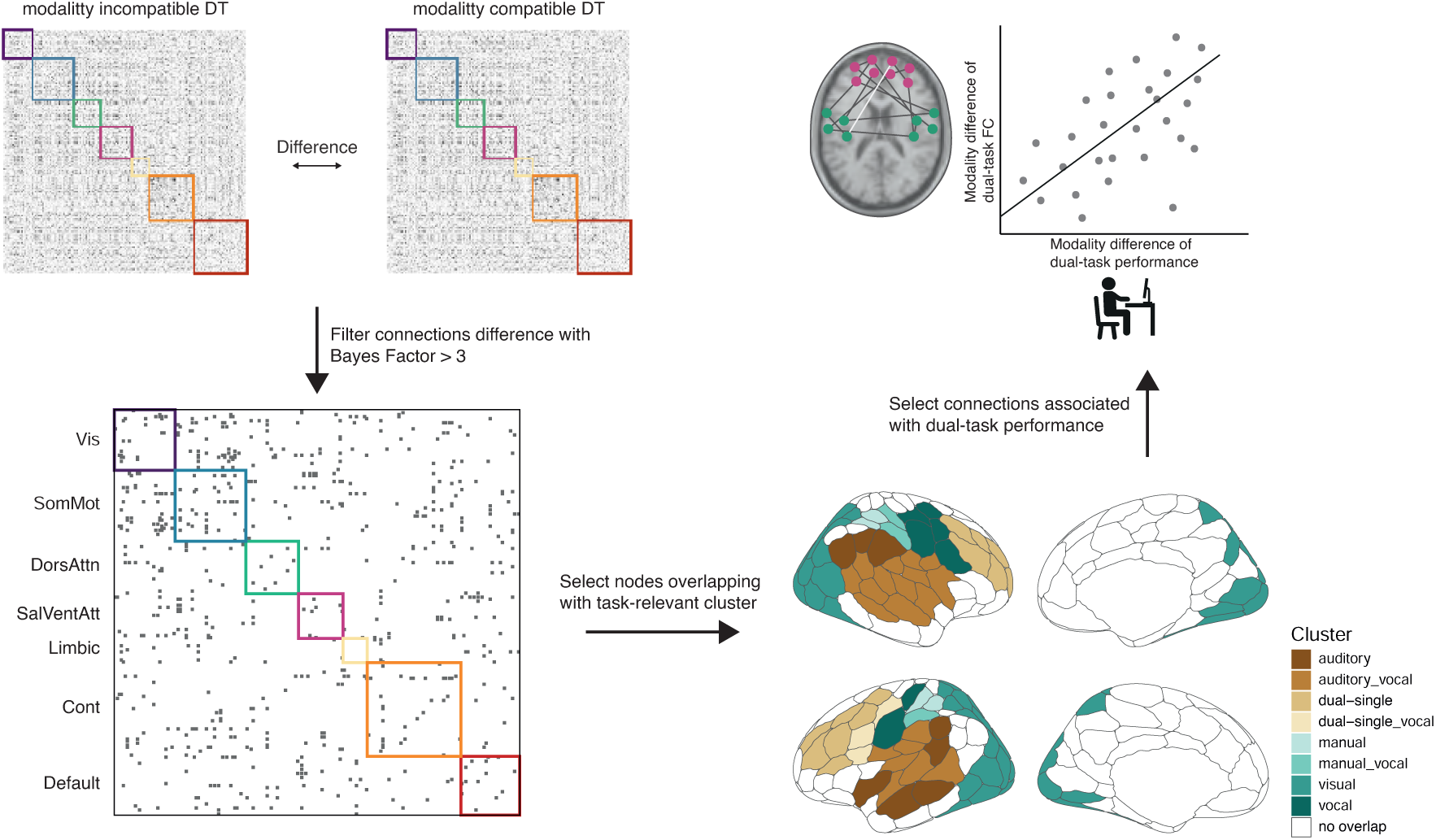
Processing steps for the second post-hoc analysis. We first calculated the difference between modality pairings of the dual-task FC matrices, next the difference was tested against zero and only connections with Bayes Factor above 3, favoring a differences remained. To relate the regions to our task-specific activity clusters we only selected regions that overlapped. Finally, we focues only on connections with a significant correlation (p < .05) between the modality difference between dual-task performance and FC difference.

Finally, to investigate which connection differences are related to the behavioral difference, we calculated partial Spearman correlation coefficients between behavioral performance (difference between modality pairings for the BIS parameter) and the FC difference between modality pairings (all connections). We controlled for differences in age, gender, and framewise displacement.

Note that for the final interpretation and discussion, we focus on the connections significantly correlated with behavior (uncorrected *p < .*05) and report the parameter *rho* for each connection.

### Data availability statement

We uploaded the BIDS transformed data (not preprocessed) to OpenNeuro: https://doi.org/10.18112/openneuro.ds005038.v1.0.1 while the beta-images, which were used to define the group-based activity clusters, are available on Neurovault: https://identifiers.org/neurovault.collection:16842 All scripts, stimulus material, and behavioral data can be accessed on OSF: https://osf.io/w9hsu/.

## Results

### Pre-registered Analysis

#### Robust difference between modality pairings in behavioral dual-task costs

We combined reaction times and accuracies into the balanced integration score (BIS) (Liesefeld & Janczyk, 2019; Mueckstein et al., 2025) to account for the dependency between the two behavioral performance parameters (see averaged reaction times and accuracies, Table S3). Please note that the behavioral data were already reported in Mueckstein et al. (2025) focusing on group-specific practice effects using a subsample of the current sample. Dual-task costs were calculated as the difference between single and dual tasks, where higher BIS scores correspond to higher dual-task costs (i.e., higher reaction times and error rates in dual tasks compared to single tasks). The modality pairings differed significantly, *t*(60) = -14.36, *p* < .001 with higher dual-task costs for the modality-incompatible pairing (*M* = 4.13, *SE* = 0.14) compared to the modality-compatible pairing (*M* = 2.03, *SE* = 0.14). The Bayes factor confirmed the strong difference as providing extreme evidence, *M* = −2.08, 95% HDI [−2.40, −1.81], BF_10_ = 1.05 × 10^18^ (see Figure 1C). This result replicates several previous findings in the field, demonstrating that performance with modality-incompatible pairings in the dual task is more error-prone and slower compared to performance with modality-compatible pairings (Göthe et al., 2016; Hazeltine et al., 2006; Mueckstein et al., 2022; Stelzel et al., 2006).

#### No difference in single-tasks FC similarity between modality pairings

To address our first neural research question, we compared the network FC similarity between modality pairings in single tasks. There was no significant effect of modality pairings in single-task FC similarity in any network combination (see Figure 4B). The highest t-value for the factor modality pairing was observed for the FC similarity within the ventral attentional network (p-value uncorrected), *t*(63) = 1.53, *p* = .132. This is confirmed by paired Bayesian t-test, which provided anecdotal to moderate evidence for the hypothesis that there is no difference between the two modality pairings (largest for the within comparison in the ventral attentional network, *M* = −0.01, 95% HDI [−0.01, 0.00], BF_10_ = 0.43 and smallest BF_10_ for between networks in the sensorimotor network, *M* = 0.00, 95% HDI [−0.01, 0.01], BF_10_ = 0.14; for detailed results and Bayes factors for all network combinations, see online material, Table S4 - S6). Figure 4A depicts the difference between modality pairings within the ventral attentional network. Thus, we observed no evidence that the single-task FC for modality-incompatible tasks (i.e., AM, VV) overlap differently than those for modality-compatible tasks (i.e., VM, AV).

**Figure 4.**
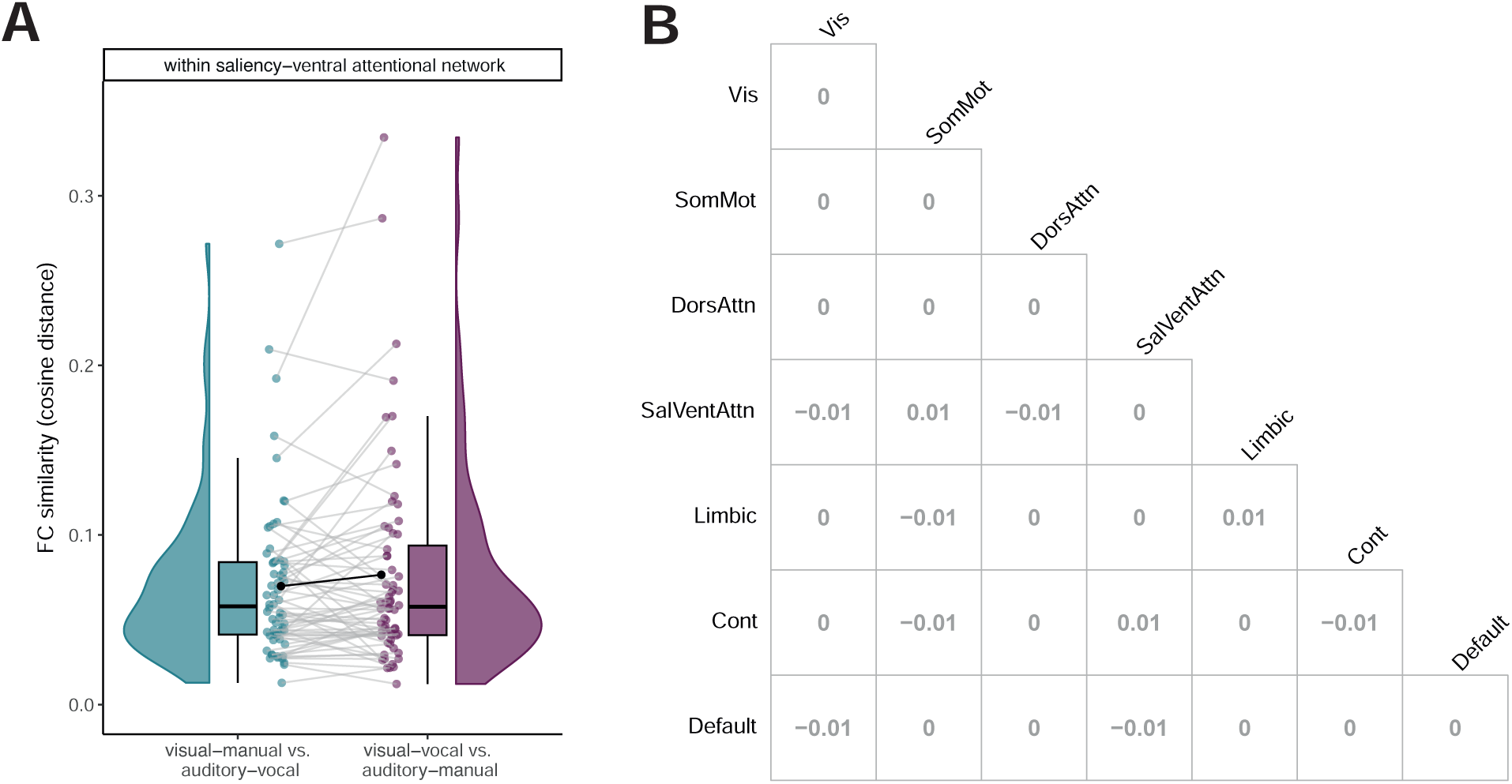
FC similarity between single task modality pairings. **A**: FC similarity within the ventral attentional network per modality pairing during single task. The graph depicts distribution, boxplot, individual data and the mean for each modality pairing of the cosine distance (not corrected for age, gender and framewise displacement). We detected no significant difference in FC similarity (cosine distance) between the two modality pairings. **B**: Matrix shows the difference between modality pairings for the FC similarity for each network combination. The diagonale of the matrix describes the within network difference. We identified no significant difference for any of the network combinations. Vis = visual network, SomMot = somato-motor network, DorsAttn = dorsal attentional network, SalVentAttn = saliency-ventral attentional network, Limbic = limbic network, Cont = control network, Default = default mode network.

**Figure 5.**
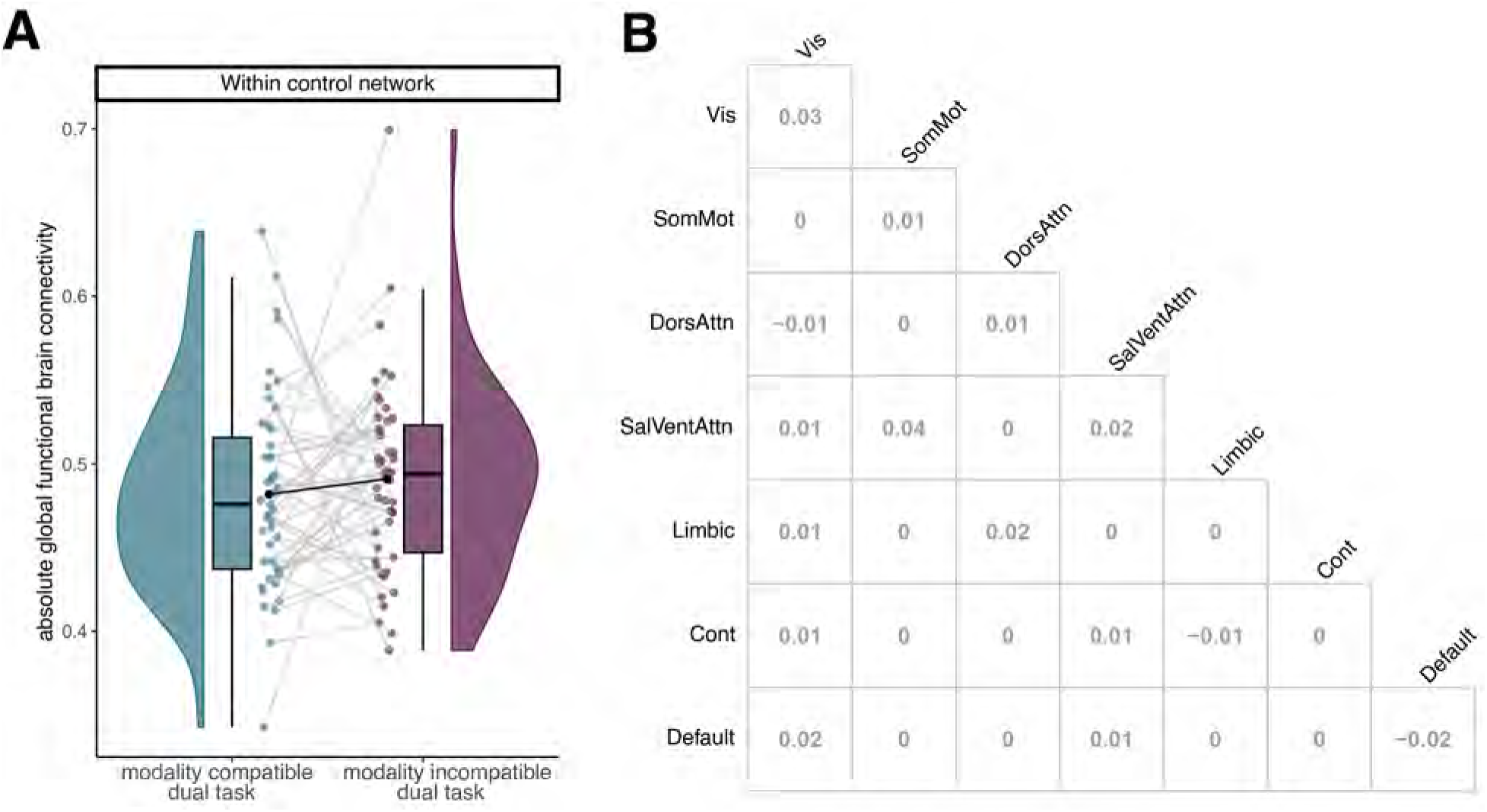
Whole-brain functional connectivty between dual-task modality pairings. **A**: Absolute whole-brain functional connectivity within the control network per modality pairing during dual task. The graph shows the distribution, boxplot, individual data and the mean for each modality pairing of the averaged absolute FC (not corrected for age, gender, framewise displacement). We found no significant difference between the dual-task modality pairings. **B**: Difference between modality pairings for the absolute whole-brain functional connectivity for each network combination. The diagonale of the matrix describes the within network difference. We found no significant difference for any of the network combinations.

#### No difference in FC strength between dual tasks

In our second research question, we further examined the involvement of the control network during dual-task performance by comparing the absolute averaged functional connectivity for each within- and between-network combination between the modality pairings. There was no significant effect of modality pairings in functional connectivity during dual-task performance in any brain network combination. The highest t-value for the factor modality pairing was observed for functional connectivity within the ventral attentional network (p-value uncorrected), *t*(46) = 1.46, *p* = .150. In line with this, Bayes factors provided anecdotal to moderate evidence for the hypothesis that there is no difference between the two modality pairings during dual-task (largest BF_10_ for within comparison in the ventral attentional network, *M* = −0.03, 95% HDI [−0.08, 0.01], BF_10_ = 0.44 and smallest for the between-network combination of the default mode network, *M* = 0.00, 95% HDI [−0.02, 0.02], BF_10_ = 0.16. For detailed results and Bayes factors for all network combinations, see the online material, Table S16 - S18. These results do not confirm the assumption of a higher connectivity strength of the control network or any other network during performance of the modality-incompatible dual task compared to the modality-compatible dual task. This null result is surprising, considering the consistent involvement of frontal regions in dual-task situations and the robust behavioral modality-compatibility effect.

We repeated both analyses with multiple variations and graph measures to rule out that our null results are specific to our preprocessing and analysis decisions (Kristanto et al., 2024). Specifically, we analyzed the data by separately changing one of the following properties: schaefer100 instead of schaefer200 parcellation (Table S22 - S27), preprocessing the data without regressing out task-related brain activity (Table S28 - S33), using Pearson correlation (Table S7 - S9) and Euclidean distance (Table S10 - S12) instead of cosine distance as FC similarity measures and the relative average of the whole-brain functional connectivity (Table S19 - S21), instead of the absolute. Also, we separately compared the brain graph’s modularity (as a measure of network segregation, compare Figure S3 and Table S34, Table S35) and global efficiency (as a measure of network communication, compare Figure S4 and Table S36, Table S37) between modality pairings for single and dual tasks to directly compare the network reconfiguration. All analyses demonstrate the same results as presented above: No difference in single-task overlap and no different involvement of the control network during dual tasks, which suggests that these global network attributes do not provide the basis for understanding the neural underpinnings of the robust behavioral modality-compatibility effect.

However, considering the robust behavioral effect of modality pairings, we further explored the data with additional analyses, comparing the FC similarity between modality pairings within each task type (within single tasks and dual tasks).

### Post-hoc Analyses

#### Significant difference between single and dual task FC similarity of modality pairings

Applying a similar approach as for the modality-compatible vs. modality-incompatible single tasks, we compared the FC similarity of modality pairings between single tasks and dual tasks. This allows us to investigate whether the FC similarity between modality pairings is different between single and dual tasks, similar to the behavioral data, where we usually don’t see a difference in single tasks but a robust difference in dual tasks. First, we averaged the single-task FC matrix per modality pairing (resulting in one FC for modality-compatible single tasks and one for modality-incompatible single tasks) to obtain the same data structure as for the dual tasks.

We then tested if the distance between the modality-compatible and the modality-incompatible FC differs between single tasks and dual tasks. We used the same model as for the single-task FC similarity analysis and compared again within and between networks.

The linear mixed-effects model revealed significant differences between single-task and dual-task FC similarity in all within and between network combinations (all *p < .*001, BH corrected), with higher values for the dual task (averaged over the between-network combinations for each network), *M* = 0.75, *SE* = 0.01, compared to the single task, *M* = 0.08, *SE* = 0.00. The smallest t-value for the factor task type was observed for the FC similarity within the dorsal attentional network, *t*(44) = −30.75, *p < .*001. Compare Figure 6 for the differences between task types for all network combinations. Bayes factors confirmed this effect, providing strong evidence for the difference between single tasks and dual tasks (paired t-test, all BF_10_ *>* 3.96 × 10^19^, for detailed results see online material, Table S13 - S15).

**Figure 6.**
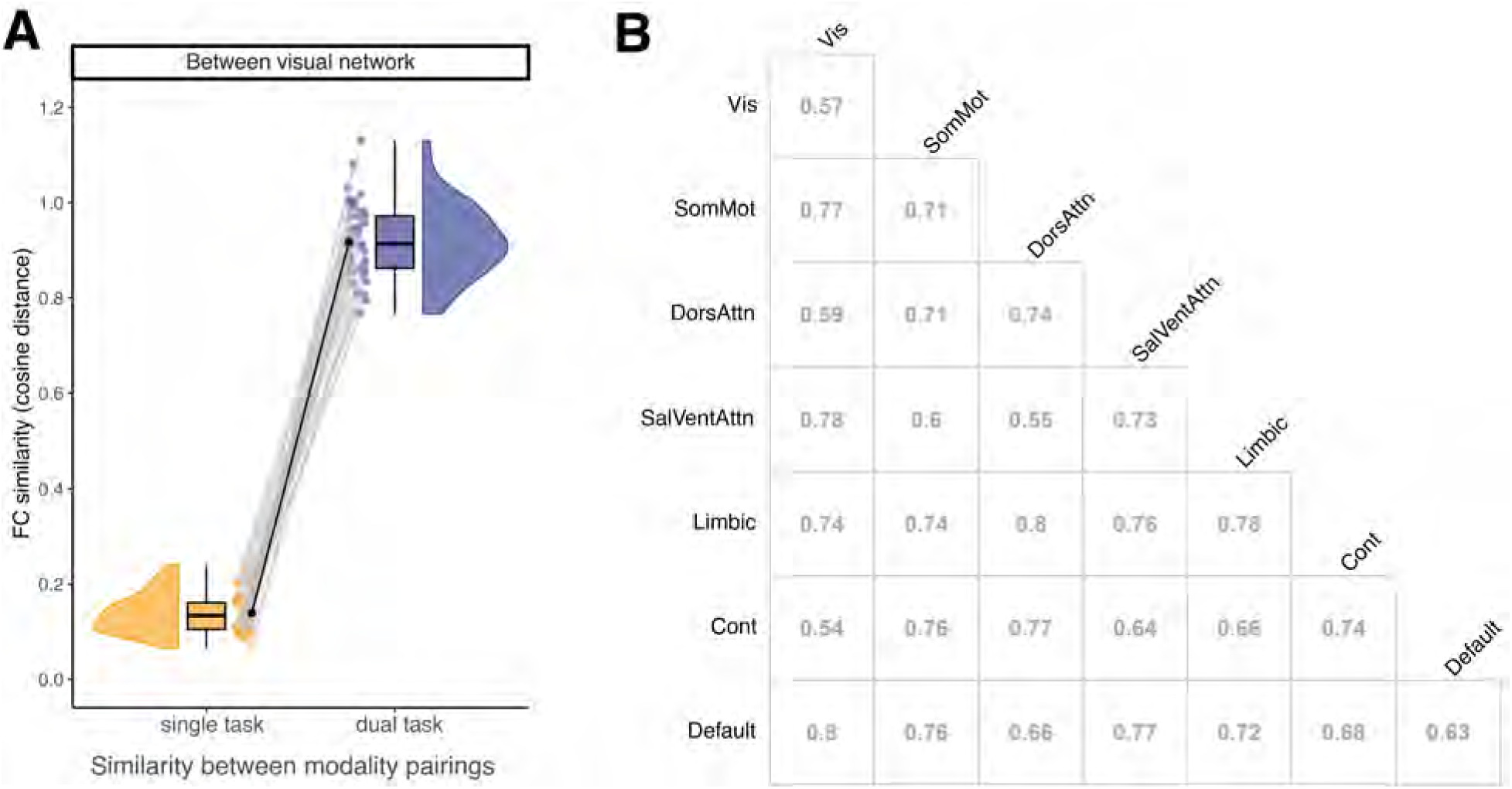
FC similarity between modality pairings for single and dual tasks. **A**: FC similarity between modality pairing between the visual network and all other networks per task type. The graph provides distribution, boxplot, individual data and the mean for each task type. We identified significant difference in FC similarity (cosine distance) of modality pairings between single and dual task. **B**: Matrix demonstrates the difference single and dual task for the cosine distance between modality-compatible and modality-incompatible for each network combination. The diagonale of the matrix describes the within network difference. We identified significant difference for all of the network combinations.

These results imply two aspects. Firstly, FC similarity of the modality pairings is significantly smaller for dual tasks than for single tasks. Secondly, when comparing the absolute values of the cosine distance, we observed values close to 0 (vectors are proportional to each other, as 1 − *cos*(0*^◦^*) = 1 − 1 = 0) for the two averaged single tasks. This indicates that the modality-compatible single-task FC is rather similar to the modality-incompatible single-task FC. In contrast, the dual task values were close to 1 (vectors are orthogonal to each other, as 1 − *cos*(90*^◦^*) = 1 − 0 = 1), suggesting a substantial distance between the modality-compatible dual task and the modality-incompatible dual task. To further understand this difference in FC similarity and, particularly, to examine which characteristics of this difference are associated with behavioral differences, we conducted an additional post-hoc analysis considering the raw dual-task FCs.

#### Characterizing the difference between dual-task functional connectivity

We calculated the difference between modality-compatible and modality-incompatible FC per participant and selected only the connections with a robust (BF_10_ *>* 3) difference between pairings during dual-task performance. Further, we focused on those brain regions from the schaefer200 parcellation, which overlap with the clusters a priori identified as task-related (univariate analysis of the localizer task), as we were interested in the neural mechanisms of the modality-compatibility effect and aimed at identifying which specific functional brain connections contributed to this effect. The remaining connections are depicted in Figure 7A and an overview of the location of each region in the task-related clusters in Figure 7B. On a descriptive level, it is remarkable that after this selection process, more interhemispheric connections between frontal regions and temporoparietal regions remain, together with highly connected regions, which overlap with the visual cluster. Both showed significantly higher FC during the modality-incompatible dual task compared to the modality-compatible dual task (purple connections). On the other hand, we observed more anterior-posterior connections with a higher FC during the modality-compatible dual task compared to the modality-incompatible dual task. To identify connections relevant for behavior, we calculated partial Spearman correlations between FC differences and behavioral difference scores between both dual tasks, controlling for age, gender, and framewise displacement (for an overview of connections significantly related to performance without the requirement that they are significantly different between modality pairings, see Figure S2). Filtering the remaining connections for their relation to dual-task performance (uncorrected *p < .*05) revealed three connections (see Figure 8A). The first connection was present between the left inferior frontal sulcus (IFS) and the right superior temporal gyrus (STG) and depicted a positive association with behavioral performance, i.e., the higher the FC during the modality-incompatible dual task compared to the modality-compatible dual task, the better the performance during the modality-incompatible dual task, *r* (45) = 0.30. Interestingly, the IFS region overlaps with the dual-task-related cluster and the STG with the auditory cluster. This correlation indicates a strong interaction between a frontal control region and a sensory auditory region that guides the successful performance of the modality-incompatible dual task, assumed to involve modality-based crosstalk. The second connection was localized within the visual cluster between the left fusiform gyrus (FFG) and the right V5. We found a negative correlation, indicating that the higher the functional connectivity during the modality-compatible dual task compared to the modality-incompatible dual task, the better the performance during the modality-incompatible dual task, *r* (45) = -0.36. The third significant connection evolved between the right superior frontal gyrus (SFG), which overlaps again with the dual-task-related cluster, and the right superior parietal lobule (SPL), which overlaps with the parietal part of the visual cluster. Similar to the IFS-STG connection, we found a positive correlation for the SFG-SPL connection, *r* (45) = 0.30.

**Figure 7.**
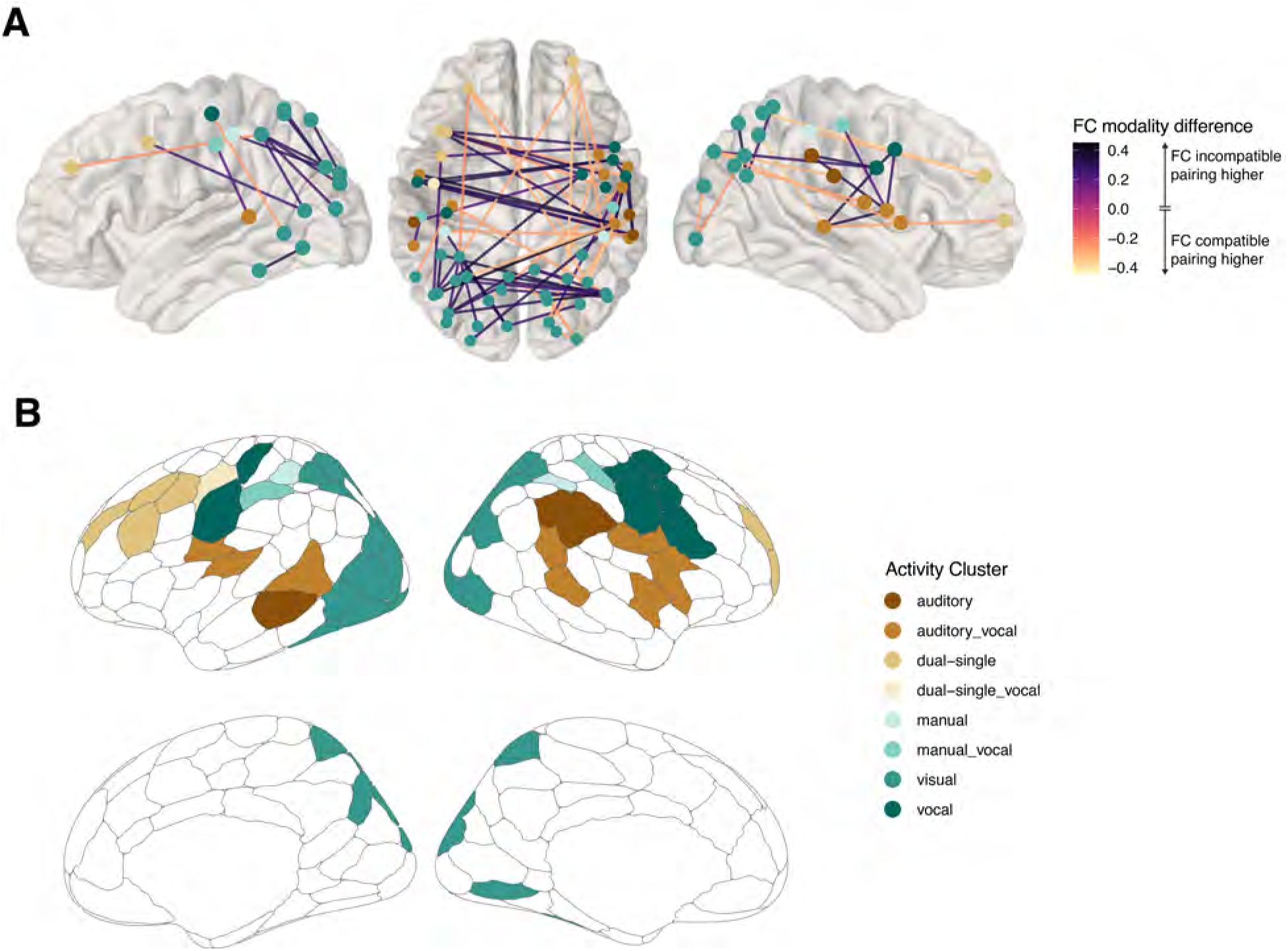
Significant different connections between modality pairings during dual tasks. **A**: The graph depicts the connections which are significantly different between the dual-task modalities, restricted to regions overlapping with task-related activity clusters. Color of regions indicate the corresponding activity cluster, color of connections the difference value of functional connectivity (FC) between modality-incompatible dual task (positive values) and modality-compatible dual task (negative values). The two sagital views contain only regions and connections within the corresponding hemisphere. The superior view contains all significant edges. **B**: The graph depicts the schaefer200 parcellation in medial and sagital view. Color of the regions indicate the corresponding task-related activity clusters with which the region overlaps. White regions did not overlap with any of our activity clusters.

**Figure 8.**
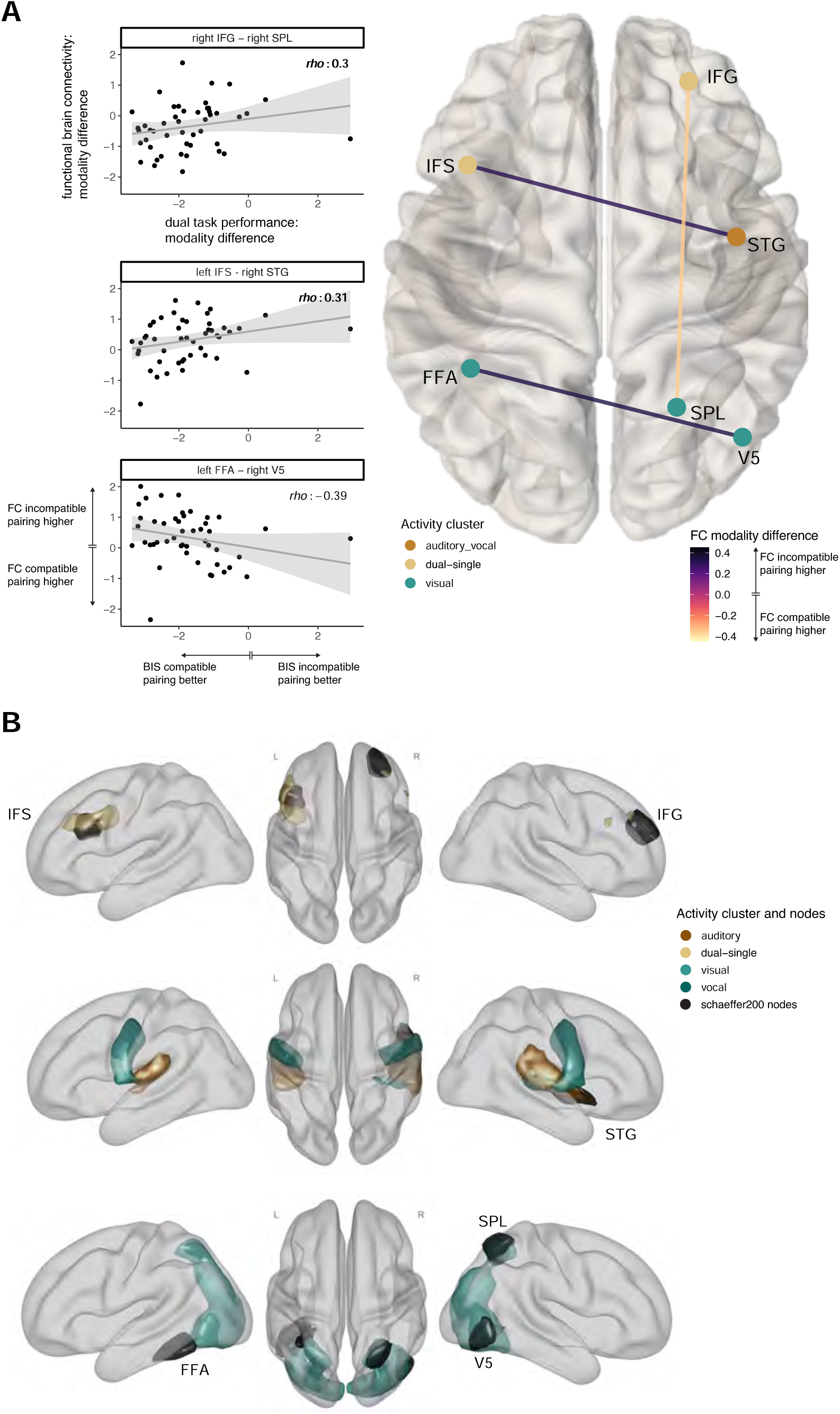
Relation between dual-task behavior and functional connectivity. **A**: The superior view of the brain with the three connections significantly different between dual-task modalities and significantly related to the dual-task behavior. Color of regions indicate the corresponding activity cluster, color of connections the difference value of functional connectivity (FC) between modality-incompatible dual task (positive values) and modality-compatible dual task (negative values). Each scatter plot corresponds to one connection and depicts each individual as one point, whereas the y-axis represents the difference score between modalities of the functional connectivity during dual tasks and the x-axis the difference score between modalities of the behavioral performance during dual task, operationalized as BIS parameter. The corresponding rho value is based on the partial correlation, corrected for age, gender and framewise displacement. **B**: Anatomical location of relevant schaefer200 regions for brain-behavior relation, together with the task-based activity clusters. Labels correspond to the Schaefer regions, colors to the activity clusters. Label abbreviations: IFG = inferior frontal gyurs, STG = superior temporal gyrus, SPL = superior parietal lobule, V5 = middle occipital gyrus, FFA = fusiform area, IFS = inferior frontal sulcus

Note, however, the difference that absolute mean functional connectivity was significantly higher during the modality-compatible dual task compared to the modality-incompatible dual task (light edge colors in Figure 8A).

Overall, these post-hoc analyses revealed significant differences on the level of individual functional brain connections related to dual-task behavior between the two dual-task pairings These were observed in regions commonly involved in multitasking scenarios and modality-based crosstalk (see Figure 8B for anatomical overlap between Schaefer regions and activity clusters).

## Discussion

While multitasking is omnipresent in everyday life, little is known about the neural basis of modality-based multitasking costs. Previous neuroimaging studies focused on univariate ROI analysis (Stelzel et al., 2006) and multivoxel pattern analysis (MVPA) (Mueckstein et al., 2025). We complement this research of modality-based crosstalk by adopting a network neuroscience perspective. Specifically, we investigated whether the single-task similarity of whole-brain functional connectivity differs between modality pairings, similar to the overlapping task representations revealed by the MVPA (Mueckstein et al., 2025). Additionally, we examined whether whole-brain connectivity during dual-task processing supports modality-dependent control network involvement.

Considering the robust differences in behavioral performances between modality-compatible and modality-incompatible pairings in dual tasks, it was surprising that we did neither find a significant difference between single-task FC similarity nor different involvement of any specific network between the modality pairings during dual tasks. However, post-hoc analyses comparing the FC similarity of modality pairings further revealed high FC similarity for the single tasks but low FC similarity for the dual tasks.

We additionally examined the differences between the dual-task FCs, focusing on predefined task-related clusters. Multiple functional brain connections differed significantly between the two modality pairings during dual-task performances, explaining the low FC similarity compared to the single tasks. Interestingly, we identified significantly higher local connectivity between frontal control and sensory regions in the auditory cortex. Individual differences in this connection were associated with behavioral differences between modality-incompatible and modality-compatible dual-task performance. This finding informs the assumed interplay of modality-based crosstalk in sensory regions and the recruitment of cognitive control to resolve it, as cognitive theories of multitasking would predict.

The fact that we did not find any whole-brain differences between modality pairings based on FC strength, FC network similarity, FC network modularity, or FC global efficiency is surprising, as many other studies reported robust differences in global network parameters for different cognitive demands. For example, Cohen and D’Esposito (2016) revealed significantly lower modularity and higher global efficiency of FC for an n-back task compared to a resting state, indicating that the transition between rest and task is accompanied by a reconfiguration of functional connectivity (Alavash et al., 2019; Cohen & D’Esposito, 2016; Cole et al., 2019; Finc et al., 2020). Similar effects were observed for the comparison between two working memory tasks (0-back vs. 2/3-back), which suggests that increased cognitive demand results in a decrease in network modularity (Braun et al., 2015; Finc et al., 2017, 2020; Gallen et al., 2023; Kaposzta et al., 2021; Krienen et al., 2014; Liang et al., 2016). This reduced segregation (or increased integration) of the functional brain network is reflected in similarity measures (such as cosine distance) (Alexander-Bloch et al., 2013). However, our results indicate no evidence for global reorganization between modality pairings. Thus, the data suggest that the modality pairings seem not to differ in their working-memory load as different n-back variations do. In contrast, despite the robust behavioral differences, the differences between modality pairings on the neural level are rather subtle. This aligns with previous research that applied a univariate group-based analysis, which did not reveal significant dual-task-related differences between modality pairings on a whole-brain level but only differences in individually defined ROIs (Stelzel et al., 2006).

The challenge in this line of research is to understand the underlying neural mechanisms responsible for the robust behavioral difference between the modality pairings in dual tasks. Considering our and previous null results, instead of expecting different mechanisms, it might also be possible that both dual-task pairings are simply processed in the same way on a neural level, resulting in different behavioral outcomes. Employing this neural processing mode for the modality-compatible dual-task works well but fails (i.e., produces higher behavioral dual-task costs) when facing the modality-incompatible pairing with additional modality-based crosstalk. These brain processes need to be reconfigured to meet the demands of the modality-incompatible pairing, which may then lead to performance improvement. The first hint that this reconfiguration process might require some practice is provided by the pre-post comparison of the current data applying MPVA (Mueckstein et al., 2025). In that paper, we demonstrated that the change in single-task representational overlap correlated with the behavioral improvement of performance after a highly specific practice intervention, specifically for the participants practicing the modality-incompatible pairing. Future studies should extend this practice approach to a larger sample to account for individual differences and focus on dual-task reconfiguration to strengthen this hypothesis. However, in other domains, there is already evidence for such a practice-related network reconfiguration, for example, for motor skills (Bassett et al., 2015) and working memory (Finc et al., 2020).

Even though we did not obtain global differences between modality pairings, differences in the strength of individual functional brain connections and their relation to dual-task performance align with previous dual-task studies (Stelzel et al., 2009) and specific modality-related research. The first of the three functional connections that differed significantly in strength between the modality-compatible and the modality-incompatible pairing links the left inferior frontal sulcus (IFS) with the right superior temporal gyrus (STG). These brain regions overlap with our task-based activity clusters of the dual-single task contrast (IFS), associated with cognitive control (Diveica et al., 2023; Worringer et al., 2019) and the auditory-vocal cluster (STG), related to auditory processing (Figure 8B). The correlation with behavioral performance indicates better performance for the modality-incompatible pairing for individuals with higher FC during the modality-incompatible dual task compared to the modality-compatible dual task. The involvement of the auditory regions is in line with our results from applying MVPA, which provided evidence that the single-task representations overlap more for the modality-incompatible pairings, specifically in the auditory regions before the practice intervention (Mueckstein et al., 2025). Generally, this temporal region seems sensitive to auditory feedback (Heinks-Maldonado et al., 2005; Tourville et al., 2008), which further strengthens our assumption of modality-based crosstalk being most prominent in the auditory regions due to the overlap between the auditory stimulus in one task and the expected auditory action-effect of vocal responses in the other task. Overall, the localization of this connection, as well as its relation to behavioral performance suggests that auditory regions need to be regulated by frontal control regions to resolve the modality-based crosstalk and thus improve performance successfully.

The second significantly different connection was observed within the visual cluster, linking the left fusiform area (FFA) with the right middle occipital gyrus (V5). This relation strengthens the assumed role of sensory regions in the emergence of modality-based crosstalk, and it might be associated with visual action effects, which are typically associated with manual actions. However, we observed no differences in overlap in the visual cluster with MVPA (Mueckstein et al., 2025). The effects in the fusiform area were against our expectations, thus requiring further investigation.

The third significantly different connection links the right inferior frontal gyrus (IFG) with the right superior parietal lobule (SPL). While the IFG overlaps again with our dual-single-task cluster and is commonly activated in dual-task and task-switching studies (Worringer et al., 2019), the SPL region overlaps with the parietal part of our visual cluster. This region was identified to be essential for response-code conflict (Paas Oliveros et al., 2023) and was demonstrated to disrupt control conflict on the perceptual level in a TMS study (Soutschek et al., 2013). In summary, both regions are often associated with different kinds of conflict requiring cognitive control. The correlation suggests that for successful behavioral performance in the modality-incompatible pairing, this functional connection needs to be strengthened during the modality-incompatible dual task compared to modality-compatible dual task.

Cognitive theories of multitasking propose that overlapping task representations increase between-task crosstalk, increase dual-task costs, and require cognitive control (Frings et al., 2020; Janczyk et al., 2014; Koch, 2009; Logan & Gordon, 2001). This overlap in task representation is not only relevant between stimuli and responses but also on the level of the anticipated effect of responses and their modality (Greenwald, 1970). Over a series of experiments (Schacherer & Hazeltine, 2020, 2021, 2023) provided compelling behavioral evidence for the importance of crosstalk between stimulus modality in one task and the anticipated action effect of the other task (modality-based crosstalk). Our neural data, analyzed with MVPA, further support these conclusions by providing evidence that overlap between neural single-task representations in auditory regions contributes essentially to the dual-task costs. The current study complements this line of research by adopting a network neuroscience perspective and by revealing that global brain reconfiguration processes are not triggered by differences in modality compatibility.

Further post-hoc analyses demonstrated that during dual-task performance, the connection between auditory regions and frontal control regions is indicative of behavioral performance. This provides additional evidence for the emergence of modality-based crosstalk. Neural network differences between modality pairings are subtle and local, undetected by global similarity or network measures. Overall, our study reveals that robust and large behavioral differences are not necessarily related to global neural effects in functional brain connectivity but can be related to local differences in individual functional brain connections.

## Acknowledgement

Grammarly was used for language editing. We thank Elisa Arnold, Friederike Glueck, Gregory Gutmann, Lea Lowak, Max Nowaczyk and Oliver Stegmann for assisting in data collection and preprocessing of the vocal data. Neuroimaging was performed at the Cognitive Center for Neuroscience Berlin and was technically supported by Christian Kainz and Till Nierhaus. This work was financially supported by the German Research Foundation, Priority Program SPP 1772 [grant numbers: STE 2226/4-2; GR 3997/4-2; HE 7464/1-2; RA 1047/4-2].

## Supplementary Material

### General information

The following abbreviations are used in the reporting of the models:

|  |  |
| --- | --- |
| STc | modality-compatible single task |
| STi | modality-incompatible single task |
| DTc | modality-compatible dual task |
| DTi | modality-incompatible dual task |
| sexM | male participants |
| meanFD | averaged framewise displacement (parameter indicating the amount of headmovement in the fMRI scanner) |
| run | indicate the number of valid runs for the single tasks (included as confound, varies between 3-8) |

Significance is depicted as stars in the following way: \*\*\*: $p < .001$ , \*\*: $p < .01$ , \*: $p < .05$ .

**Table S2.** Sample size and exclusion reasons per analysis.

|  | Excluded<br>participants | Exclusion reason |
| --- | --- | --- |
| Behavioral analysis (N=61) | 10 | error rate |
| Localizer task (N = 53) | 10 | error rate |
|  | 7 | response-modality error |
|  | 1 | head movement |
| Connectivity single task (N=64) | 7 | head movement |
| Connectivity dual task (N=47) | 10 | error rate |
|  | 14 | head movement |
| Connectivity single vs. dual task (N=45) | 10 | error rate dual task |
|  | 14 | head movement dual task |
|  | 2 | head movement single task |

Showing numbers of excluded participants. Note the sample of the 'Connectivity single task' analysis was used as well for the decoding analysis at Pre timepoint reported in Mueckstein, et al., 2025. In the decoding analysis, five more participants could be included as the head-movement criteria were less restrictive for the MVPA compared to the connectivity analysis.

**Figure S1.**
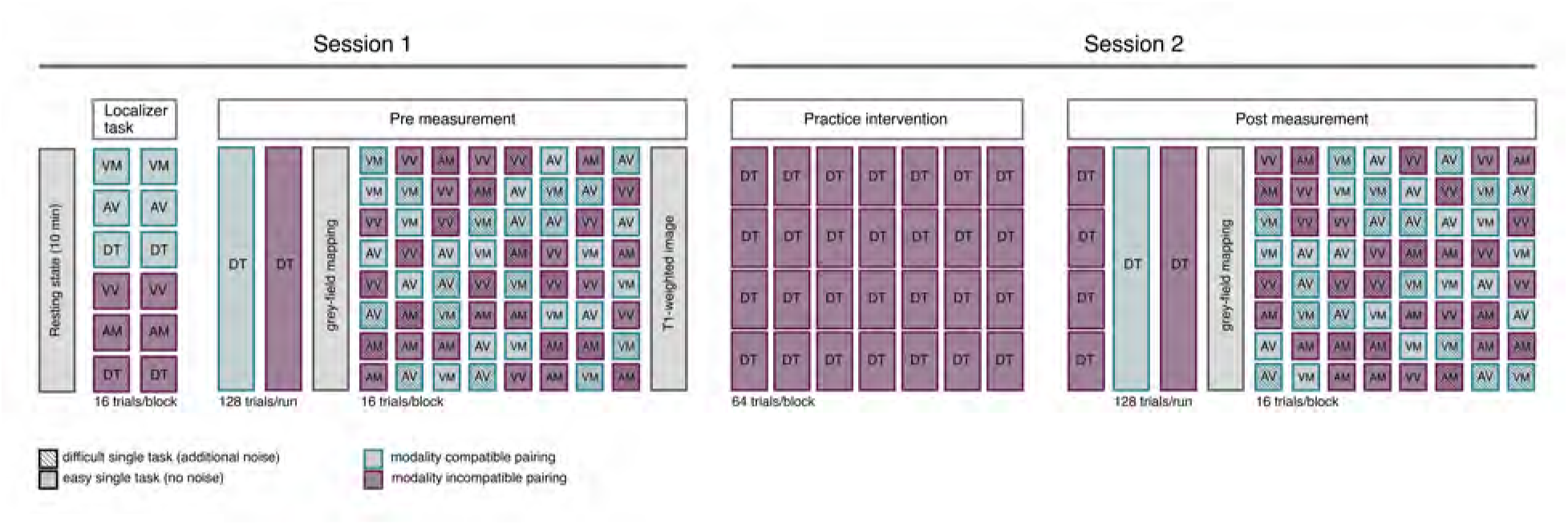
Structure of the fMRI Pre/Post session with a block design. The order of blocks in the Localizer Task was fixed to single task, dual task per modality mapping. The modality of the first task (visual or auditory) and the order of the modality pairing was balanced across participants. The order of the two dual-task runs in the pre session was balanced across participants as well. The following eight runs contained only single-task runs. Each run contained two blocks for the four different single tasks, one per task difficulty level, in randomized order. The practice intervention was completed outside the scanner on a computer screen, depending on the group-assignment participants either worked on modality-compatible or modality-incompatible dual tasks. VM = visual-manual, AV = auditory-vocal, VV = visual-vocal, AM = auditory-vocal, DT = dual task

**Table S3.**
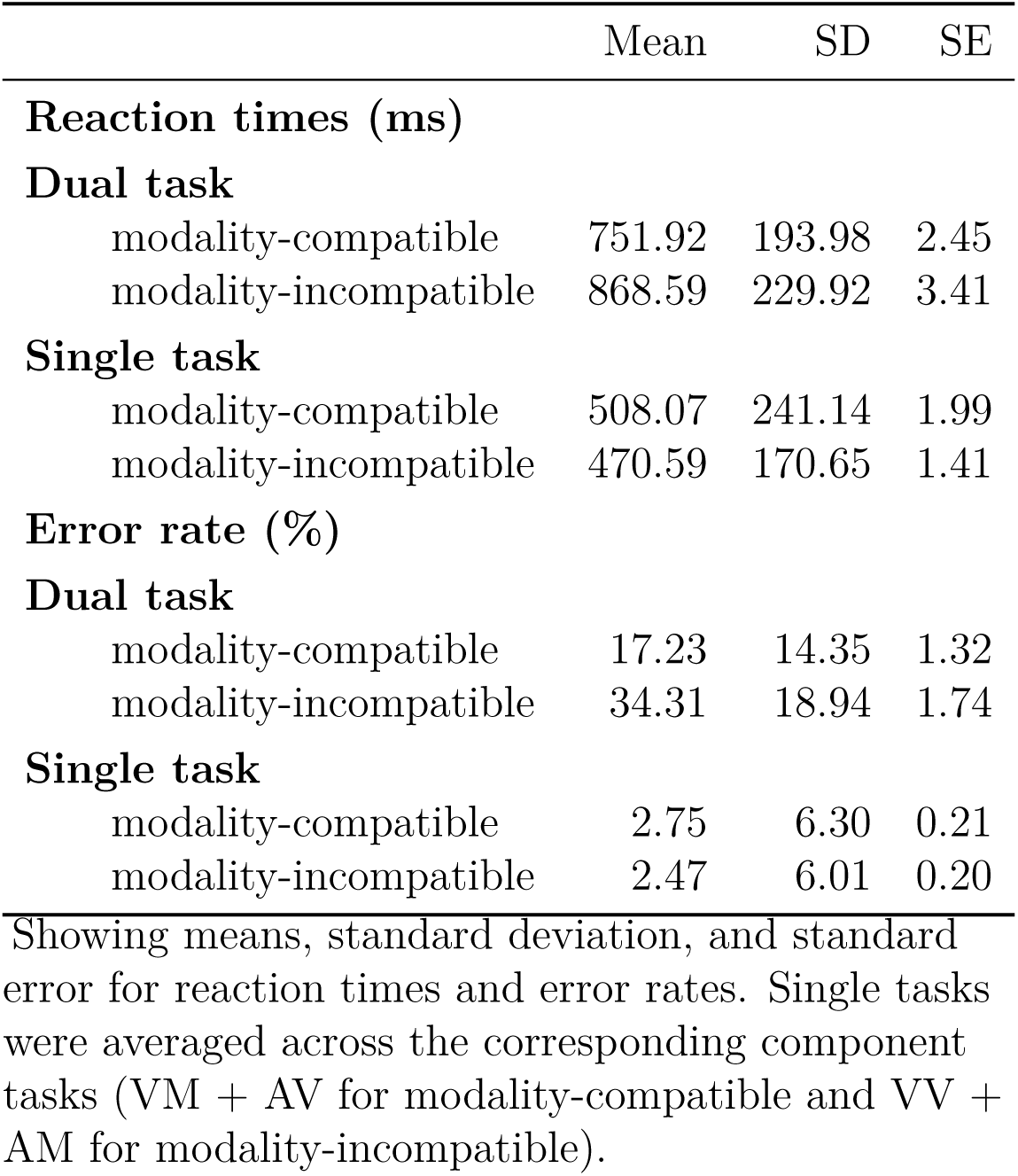
Reaction times and error rates per task type, and modality mapping.

|  | Mean | SD | SE |
| --- | --- | --- | --- |
| **Reaction times (ms)** |  |  |  |
| **Dual task** |  |  |  |
| modality-compatible | 751.92 | 193.98 | 2.45 |
| modality-incompatible | 868.59 | 229.92 | 3.41 |
| **Single task** |  |  |  |
| modality-compatible | 508.07 | 241.14 | 1.99 |
| modality-incompatible | 470.59 | 170.65 | 1.41 |
| **Error rate (%)** |  |  |  |
| **Dual task** |  |  |  |
| modality-compatible | 17.23 | 14.35 | 1.32 |
| modality-incompatible | 34.31 | 18.94 | 1.74 |
| **Single task** |  |  |  |
| modality-compatible | 2.75 | 6.30 | 0.21 |
| modality-incompatible | 2.47 | 6.01 | 0.20 |

Showing means, standard deviation, and standard error for reaction times and error rates. Single tasks were averaged across the corresponding component tasks (VM + AV for modality-compatible and VV + AM for modality-incompatible).

### FC similarity between single tasks

**Table S4.** FC similarity (cosine distance) between single tasks per between-network combination, full model results.

|  |  | value | std. error | df | t-value | p-value |  |
| --- | --- | --- | --- | --- | --- | --- | --- |
| control | stateSTi | 0.000 | 0.005 | 63 | 0.081 | 0.936 |  |
|  | meanFD | 0.279 | 0.235 | 59 | 1.184 | 0.241 |  |
|  | run | -0.019 | 0.004 | 59 | -4.691 | 0.000 | *** |
|  | age | -0.002 | 0.002 | 59 | -0.842 | 0.403 |  |
|  | sexM | -0.009 | 0.011 | 59 | -0.765 | 0.447 |  |
| default | stateSTi | -0.002 | 0.004 | 63 | -0.390 | 0.698 |  |
|  | meanFD | 0.457 | 0.228 | 59 | 2.003 | 0.050 | * |
|  | run | -0.017 | 0.004 | 59 | -4.325 | 0.000 | *** |
|  | age | -0.001 | 0.002 | 59 | -0.678 | 0.501 |  |
|  | sexM | -0.006 | 0.011 | 59 | -0.498 | 0.621 |  |
| dorsAtt | stateSTi | -0.003 | 0.006 | 63 | -0.511 | 0.611 |  |
|  | meanFD | 0.485 | 0.232 | 59 | 2.089 | 0.041 | * |
|  | run | -0.019 | 0.004 | 59 | -4.687 | 0.000 | *** |
|  | age | -0.001 | 0.002 | 59 | -0.365 | 0.717 |  |
|  | sexM | -0.007 | 0.011 | 59 | -0.648 | 0.520 |  |
| limbic | stateSTi | -0.007 | 0.008 | 63 | -0.842 | 0.403 |  |
|  | meanFD | 0.661 | 0.318 | 59 | 2.080 | 0.042 | * |
|  | run | -0.012 | 0.006 | 59 | -2.070 | 0.043 | * |
|  | age | -0.003 | 0.003 | 59 | -1.148 | 0.256 |  |
|  | sexM | 0.002 | 0.015 | 59 | 0.150 | 0.882 |  |
| somatomotor | stateSTi | 0.000 | 0.006 | 63 | -0.069 | 0.945 |  |
|  | meanFD | 0.344 | 0.274 | 59 | 1.255 | 0.214 |  |
|  | run | -0.014 | 0.005 | 59 | -2.932 | 0.005 | ** |
|  | age | -0.003 | 0.002 | 59 | -1.404 | 0.166 |  |
|  | sexM | -0.014 | 0.013 | 59 | -1.078 | 0.285 |  |
| ventrAtt | stateSTi | -0.001 | 0.006 | 63 | -0.162 | 0.872 |  |
|  | meanFD | 0.296 | 0.245 | 59 | 1.208 | 0.232 |  |
|  | run | -0.018 | 0.004 | 59 | -4.197 | 0.000 | *** |
|  | age | -0.002 | 0.002 | 59 | -0.927 | 0.357 |  |
|  | sexM | -0.009 | 0.012 | 59 | -0.747 | 0.458 |  |
| visual | stateSTi | -0.008 | 0.007 | 63 | -1.229 | 0.224 |  |
|  | meanFD | 0.265 | 0.279 | 59 | 0.953 | 0.345 |  |
|  | run | -0.013 | 0.005 | 59 | -2.578 | 0.012 | * |
|  | age | -0.004 | 0.002 | 59 | -1.746 | 0.086 |  |
|  | sexM | -0.012 | 0.014 | 59 | -0.895 | 0.375 |  |

line per between-network combination (e.g., connections from visual to all other networks): cosine distance = STc vs. STi + confounds. Between connections averaged for each network. Task-activity regressed out, schaefer200 parcellation

**Table S5.** FC similarity (cosine distance) between single tasks per within-network combination, full model results.

|  |  | value | std. error | df | t-value | p-value |  |
| --- | --- | --- | --- | --- | --- | --- | --- |
| control | stateSTi | 0.008 | 0.005 | 63 | 1.516 | 0.134 |  |
|  | meanFD | 0.154 | 0.177 | 59 | 0.869 | 0.388 |  |
|  | run | -0.021 | 0.003 | 59 | -6.701 | 0.000 | *** |
|  | age | 0.001 | 0.001 | 59 | 0.867 | 0.389 |  |
|  | sexM | 0.000 | 0.009 | 59 | 0.013 | 0.989 |  |
| default | stateSTi | -0.003 | 0.003 | 63 | -1.239 | 0.220 |  |
|  | meanFD | 0.406 | 0.203 | 59 | 1.997 | 0.050 |  |
|  | run | -0.021 | 0.004 | 59 | -5.960 | 0.000 | *** |
|  | age | 0.000 | 0.002 | 59 | -0.184 | 0.854 |  |
|  | sexM | 0.012 | 0.010 | 59 | 1.237 | 0.221 |  |
| dorsAtt | stateSTi | 0.001 | 0.006 | 63 | 0.175 | 0.862 |  |
|  | meanFD | 0.257 | 0.193 | 59 | 1.328 | 0.189 |  |
|  | run | -0.020 | 0.003 | 59 | -5.859 | 0.000 | *** |
|  | age | 0.000 | 0.002 | 59 | -0.009 | 0.993 |  |
|  | sexM | 0.002 | 0.009 | 59 | 0.266 | 0.791 |  |
| limbic | stateSTi | -0.005 | 0.007 | 63 | -0.791 | 0.432 |  |
|  | meanFD | 0.217 | 0.220 | 59 | 0.984 | 0.329 |  |
|  | run | -0.020 | 0.004 | 59 | -5.268 | 0.000 | *** |
|  | age | -0.001 | 0.002 | 59 | -0.684 | 0.497 |  |
|  | sexM | 0.017 | 0.011 | 59 | 1.543 | 0.128 |  |
| somatomotor | stateSTi | 0.004 | 0.004 | 63 | 1.066 | 0.291 |  |
|  | meanFD | 0.285 | 0.174 | 59 | 1.638 | 0.107 |  |
|  | run | -0.015 | 0.003 | 59 | -4.774 | 0.000 | *** |
|  | age | -0.002 | 0.001 | 59 | -1.069 | 0.289 |  |
|  | sexM | -0.002 | 0.008 | 59 | -0.242 | 0.810 |  |
| ventrAtt | stateSTi | 0.007 | 0.004 | 63 | 1.526 | 0.132 |  |
|  | meanFD | 0.244 | 0.220 | 59 | 1.111 | 0.271 |  |
|  | run | -0.014 | 0.004 | 59 | -3.656 | 0.001 | *** |
|  | age | -0.001 | 0.002 | 59 | -0.439 | 0.662 |  |
|  | sexM | 0.008 | 0.011 | 59 | 0.782 | 0.437 |  |
| visual | stateSTi | 0.002 | 0.003 | 63 | 0.621 | 0.537 |  |
|  | meanFD | 0.119 | 0.133 | 59 | 0.893 | 0.375 |  |
|  | run | -0.015 | 0.002 | 59 | -6.247 | 0.000 | *** |
|  | age | 0.001 | 0.001 | 59 | 0.650 | 0.518 |  |
|  | sexM | -0.001 | 0.006 | 59 | -0.137 | 0.892 |  |

line per within-network combination (e.g., connections within the visual network): cosine distance = STc vs. STi + confounds Task-activity regressed out, schaefer200 parcellation

**Table S6.**
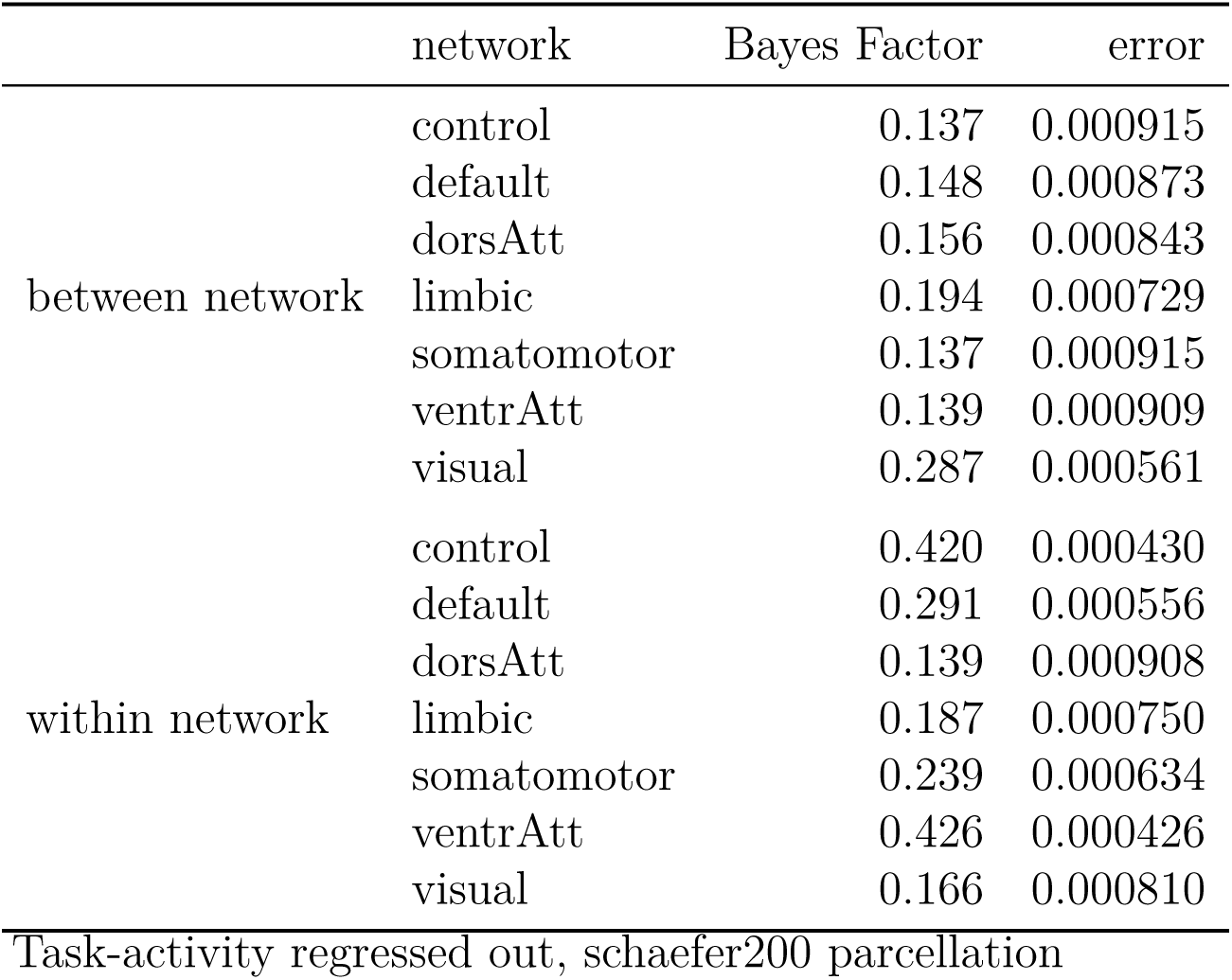
Bayes paired t-test between FC similarity (cosine distance) of the single task modalitiy pairings.

|  | network | Bayes Factor | error |
| --- | --- | --- | --- |
| between network | control | 0.137 | 0.000915 |
|  | default | 0.148 | 0.000873 |
|  | dorsAtt | 0.156 | 0.000843 |
|  | limbic | 0.194 | 0.000729 |
|  | somatomotor | 0.137 | 0.000915 |
|  | ventrAtt | 0.139 | 0.000909 |
|  | visual | 0.287 | 0.000561 |
| within network | control | 0.420 | 0.000430 |
|  | default | 0.291 | 0.000556 |
|  | dorsAtt | 0.139 | 0.000908 |
|  | limbic | 0.187 | 0.000750 |
|  | somatomotor | 0.239 | 0.000634 |
|  | ventrAtt | 0.426 | 0.000426 |
|  | visual | 0.166 | 0.000810 |
| Task-activity regressed out, schaefer200 parcellation |  |  |  |

### 0.1 Alternative FC similarity parameter

**Table S7.** FC similarity (correlation) between single tasks per between-network combination, full model results.

|  |  | value | std. error | df | t-value | p-value |  |
| --- | --- | --- | --- | --- | --- | --- | --- |
| control | stateSTIv - STIa | -0.002 | 0.005 | 63 | -0.469 | 0.641 |  |
|  | meanFD | -0.222 | 0.228 | 59 | -0.975 | 0.333 |  |
|  | run | 0.019 | 0.004 | 59 | 4.693 | 0.000 | *** |
|  | age | 0.002 | 0.002 | 59 | 0.962 | 0.340 |  |
|  | sexM | 0.012 | 0.011 | 59 | 1.056 | 0.295 |  |
| default | stateSTIv - STIa | 0.003 | 0.005 | 63 | 0.605 | 0.547 |  |
|  | meanFD | -0.319 | 0.214 | 59 | -1.491 | 0.141 |  |
|  | run | 0.018 | 0.004 | 59 | 4.740 | 0.000 | *** |
|  | age | 0.002 | 0.002 | 59 | 1.132 | 0.262 |  |
|  | sexM | 0.010 | 0.010 | 59 | 0.947 | 0.347 |  |
| dorsAtt | stateSTIv - STIa | -0.001 | 0.006 | 63 | -0.232 | 0.817 |  |
|  | meanFD | -0.392 | 0.220 | 59 | -1.783 | 0.080 |  |
|  | run | 0.020 | 0.004 | 59 | 5.062 | 0.000 | *** |
|  | age | 0.002 | 0.002 | 59 | 0.984 | 0.329 |  |
|  | sexM | 0.009 | 0.011 | 59 | 0.857 | 0.395 |  |
| limbic | stateSTIv - STIa | 0.000 | 0.007 | 63 | 0.065 | 0.948 |  |
|  | meanFD | -0.605 | 0.286 | 59 | -2.111 | 0.039 | * |
|  | run | 0.011 | 0.005 | 59 | 2.115 | 0.039 | * |
|  | age | 0.004 | 0.002 | 59 | 1.803 | 0.076 |  |
|  | sexM | 0.000 | 0.014 | 59 | 0.013 | 0.990 |  |
| somatomotor | stateSTIv - STIa | 0.001 | 0.006 | 63 | 0.092 | 0.927 |  |
|  | meanFD | -0.344 | 0.252 | 59 | -1.366 | 0.177 |  |
|  | run | 0.013 | 0.004 | 59 | 2.946 | 0.005 | ** |
|  | age | 0.003 | 0.002 | 59 | 1.553 | 0.126 |  |
|  | sexM | 0.018 | 0.012 | 59 | 1.468 | 0.147 |  |
| ventrAtt | stateSTIv - STIa | -0.001 | 0.006 | 63 | -0.215 | 0.830 |  |
|  | meanFD | -0.296 | 0.227 | 59 | -1.305 | 0.197 |  |
|  | run | 0.018 | 0.004 | 59 | 4.605 | 0.000 | *** |
|  | age | 0.002 | 0.002 | 59 | 0.979 | 0.332 |  |
|  | sexM | 0.007 | 0.011 | 59 | 0.615 | 0.541 |  |
| visual | stateSTIv - STIa | 0.006 | 0.006 | 63 | 1.021 | 0.311 |  |
|  | meanFD | -0.244 | 0.258 | 59 | -0.946 | 0.348 |  |
|  | run | 0.011 | 0.005 | 59 | 2.510 | 0.015 | * |
|  | age | 0.004 | 0.002 | 59 | 1.735 | 0.088 |  |
|  | sexM | 0.013 | 0.013 | 59 | 1.058 | 0.294 |  |

lme per between-network combination: correlation = STc vs. STi + confounds. Build-in Matlab function \*corr\* was used to calculate the Pearson correlation. Between connections averaged for each network. Task-activity regressed out, schaefer200 parcellation

**Table S8.**
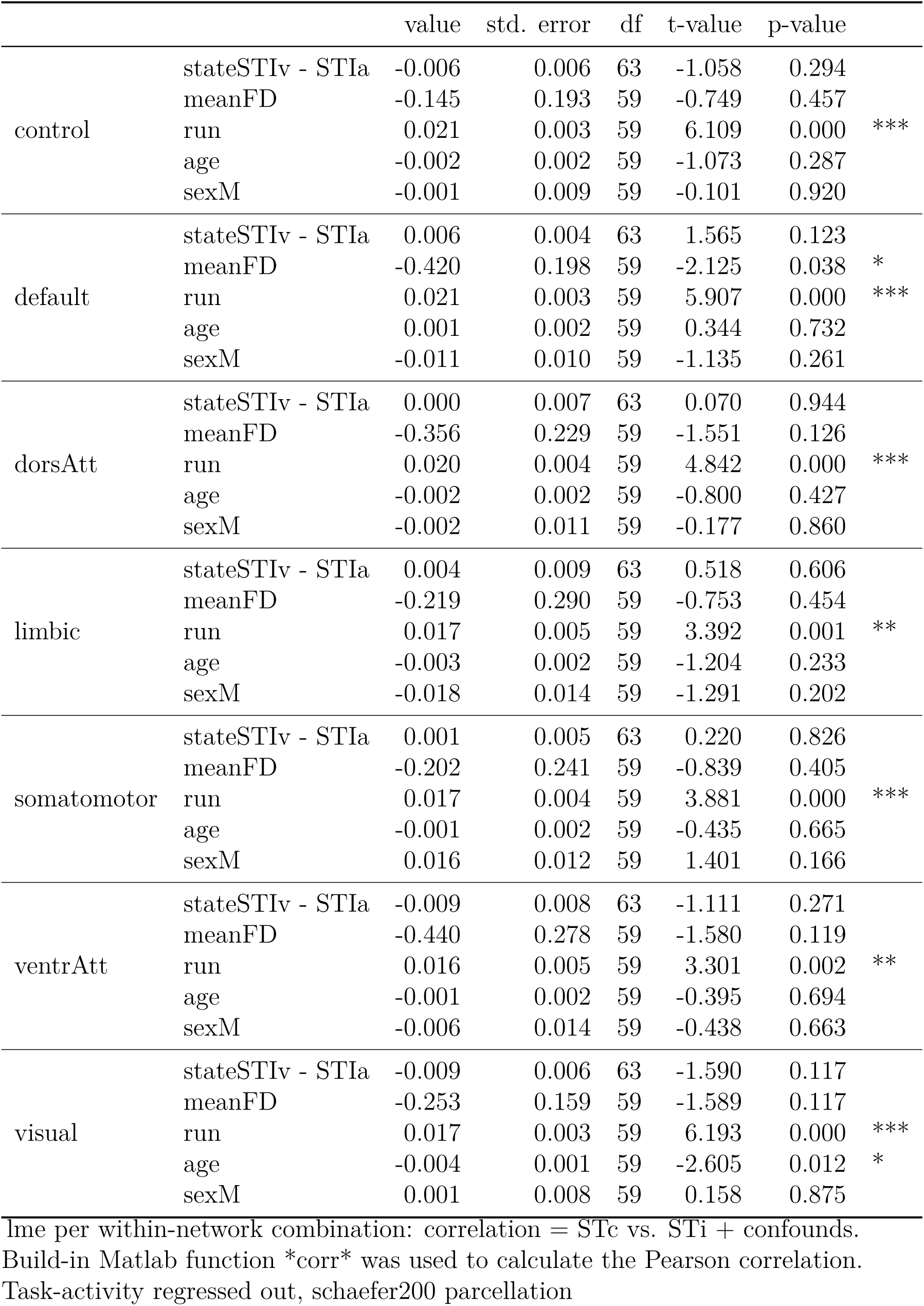
FC similarity (correlation) between single tasks per within-network combination, full model results.

|  |  | value | std. error | df | t-value | p-value |  |
| --- | --- | --- | --- | --- | --- | --- | --- |
| control | stateSTIv - STIa | -0.006 | 0.006 | 63 | -1.058 | 0.294 |  |
|  | meanFD | -0.145 | 0.193 | 59 | -0.749 | 0.457 |  |
|  | run | 0.021 | 0.003 | 59 | 6.109 | 0.000 | *** |
|  | age | -0.002 | 0.002 | 59 | -1.073 | 0.287 |  |
|  | sexM | -0.001 | 0.009 | 59 | -0.101 | 0.920 |  |
| default | stateSTIv - STIa | 0.006 | 0.004 | 63 | 1.565 | 0.123 |  |
|  | meanFD | -0.420 | 0.198 | 59 | -2.125 | 0.038 | * |
|  | run | 0.021 | 0.003 | 59 | 5.907 | 0.000 | *** |
|  | age | 0.001 | 0.002 | 59 | 0.344 | 0.732 |  |
|  | sexM | -0.011 | 0.010 | 59 | -1.135 | 0.261 |  |
| dorsAtt | stateSTIv - STIa | 0.000 | 0.007 | 63 | 0.070 | 0.944 |  |
|  | meanFD | -0.356 | 0.229 | 59 | -1.551 | 0.126 |  |
|  | run | 0.020 | 0.004 | 59 | 4.842 | 0.000 | *** |
|  | age | -0.002 | 0.002 | 59 | -0.800 | 0.427 |  |
|  | sexM | -0.002 | 0.011 | 59 | -0.177 | 0.860 |  |
| limbic | stateSTIv - STIa | 0.004 | 0.009 | 63 | 0.518 | 0.606 |  |
|  | meanFD | -0.219 | 0.290 | 59 | -0.753 | 0.454 |  |
|  | run | 0.017 | 0.005 | 59 | 3.392 | 0.001 | ** |
|  | age | -0.003 | 0.002 | 59 | -1.204 | 0.233 |  |
|  | sexM | -0.018 | 0.014 | 59 | -1.291 | 0.202 |  |
| somatomotor | stateSTIv - STIa | 0.001 | 0.005 | 63 | 0.220 | 0.826 |  |
|  | meanFD | -0.202 | 0.241 | 59 | -0.839 | 0.405 |  |
|  | run | 0.017 | 0.004 | 59 | 3.881 | 0.000 | *** |
|  | age | -0.001 | 0.002 | 59 | -0.435 | 0.665 |  |
|  | sexM | 0.016 | 0.012 | 59 | 1.401 | 0.166 |  |
| ventrAtt | stateSTIv - STIa | -0.009 | 0.008 | 63 | -1.111 | 0.271 |  |
|  | meanFD | -0.440 | 0.278 | 59 | -1.580 | 0.119 |  |
|  | run | 0.016 | 0.005 | 59 | 3.301 | 0.002 | ** |
|  | age | -0.001 | 0.002 | 59 | -0.395 | 0.694 |  |
|  | sexM | -0.006 | 0.014 | 59 | -0.438 | 0.663 |  |
| visual | stateSTIv - STIa | -0.009 | 0.006 | 63 | -1.590 | 0.117 |  |
|  | meanFD | -0.253 | 0.159 | 59 | -1.589 | 0.117 |  |
|  | run | 0.017 | 0.003 | 59 | 6.193 | 0.000 | *** |
|  | age | -0.004 | 0.001 | 59 | -2.605 | 0.012 | * |
|  | sexM | 0.001 | 0.008 | 59 | 0.158 | 0.875 |  |

lme per within-network combination: correlation = STc vs. STi + confounds. Build-in Matlab function \*corr\* was used to calculate the Pearson correlation. Task-activity regressed out, schaefer200 parcellation

**Table S9.** Bayes paired t-test between FC similarity (correlation) of the single task modalitiy pairings.

|  | network | Bayes Factor | error |
| --- | --- | --- | --- |
| between network | control | 0.153 | 0.000854 |
|  | default | 0.164 | 0.000815 |
|  | dorsAtt | 0.141 | 0.000901 |
|  | limbic | 0.137 | 0.000915 |
|  | somatomotor | 0.138 | 0.000914 |
|  | ventrAtt | 0.140 | 0.000903 |
|  | visual | 0.229 | 0.000654 |
| within network | control | 0.237 | 0.000638 |
|  | default | 0.451 | 0.000409 |
|  | dorsAtt | 0.137 | 0.000915 |
|  | limbic | 0.156 | 0.000841 |
|  | somatomotor | 0.140 | 0.000903 |
|  | ventrAtt | 0.251 | 0.000614 |
|  | visual | 0.469 | 0.000398 |
| Task-activity regressed out, schaefer200 parcellation |  |  |  |

**Table S10.** FC similarity (euclidean distance) between single tasks per between-network combination, full model results.

|  |  | value | std. error | df | t-value | p-value |  |
| --- | --- | --- | --- | --- | --- | --- | --- |
| control | stateSTIv - STIa | -0.020 | 0.098 | 63 | -0.203 | 0.840 |  |
|  | meanFD | 5.403 | 2.953 | 59 | 1.829 | 0.072 |  |
|  | run | -0.836 | 0.052 | 59 | -16.044 | 0.000 | *** |
|  | age | -0.021 | 0.025 | 59 | -0.851 | 0.398 |  |
|  | sexM | -0.001 | 0.144 | 59 | -0.004 | 0.997 |  |
| default | stateSTIv - STIa | -0.104 | 0.109 | 63 | -0.961 | 0.340 |  |
|  | meanFD | 8.859 | 3.660 | 59 | 2.421 | 0.019 | * |
|  | run | -0.990 | 0.065 | 59 | -15.331 | 0.000 | *** |
|  | age | -0.038 | 0.031 | 59 | -1.232 | 0.223 |  |
|  | sexM | -0.008 | 0.178 | 59 | -0.045 | 0.964 |  |
| dorsAtt | stateSTIv - STIa | -0.072 | 0.103 | 63 | -0.697 | 0.488 |  |
|  | meanFD | 6.855 | 3.059 | 59 | 2.241 | 0.029 | * |
|  | run | -0.786 | 0.054 | 59 | -14.560 | 0.000 | *** |
|  | age | -0.021 | 0.026 | 59 | -0.800 | 0.427 |  |
|  | sexM | -0.044 | 0.149 | 59 | -0.293 | 0.771 |  |
| limbic | stateSTIv - STIa | -0.050 | 0.069 | 63 | -0.729 | 0.469 |  |
|  | meanFD | 5.140 | 1.984 | 59 | 2.591 | 0.012 | * |
|  | run | -0.553 | 0.035 | 59 | -15.787 | 0.000 | *** |
|  | age | -0.036 | 0.017 | 59 | -2.141 | 0.036 | * |
|  | sexM | 0.098 | 0.097 | 59 | 1.013 | 0.315 |  |
| somatomotor | stateSTIv - STIa | -0.037 | 0.112 | 63 | -0.332 | 0.741 |  |
|  | meanFD | 7.849 | 3.455 | 59 | 2.272 | 0.027 | * |
|  | run | -0.917 | 0.061 | 59 | -15.045 | 0.000 | *** |
|  | age | -0.046 | 0.029 | 59 | -1.571 | 0.122 |  |
|  | sexM | 0.007 | 0.168 | 59 | 0.042 | 0.967 |  |
| ventrAtt | stateSTIv - STIa | -0.064 | 0.094 | 63 | -0.679 | 0.500 |  |
|  | meanFD | 5.438 | 2.664 | 59 | 2.042 | 0.046 | * |
|  | run | -0.748 | 0.047 | 59 | -15.912 | 0.000 | *** |
|  | age | -0.025 | 0.023 | 59 | -1.117 | 0.269 |  |
|  | sexM | -0.043 | 0.130 | 59 | -0.330 | 0.743 |  |
| visual | stateSTIv - STIa | -0.049 | 0.104 | 63 | -0.470 | 0.640 |  |
|  | meanFD | 5.107 | 2.925 | 59 | 1.746 | 0.086 |  |
|  | run | -0.847 | 0.052 | 59 | -16.408 | 0.000 | *** |
|  | age | -0.031 | 0.025 | 59 | -1.235 | 0.222 |  |
|  | sexM | 0.014 | 0.142 | 59 | 0.097 | 0.923 |  |

line per between-network combination: euclidean distance = STc vs. STi + confounds. Build-in Matlab function \*pdist\* with the parameter 'euclidean' was used to calculate the Euclidean distance. Between connections averaged for each network. Task-activity regressed out, schaefer200 parcellation

**Table S11.** FC similarity (euclidean distance) between single tasks per within-network combination, full model results.

|  |  | value | std. error | df | t-value | p-value |  |
| --- | --- | --- | --- | --- | --- | --- | --- |
| control | stateSTIv - STIa | 0.094 | 0.106 | 63 | 0.892 | 0.376 |  |
|  | meanFD | 2.565 | 2.400 | 59 | 1.069 | 0.289 |  |
|  | run | -0.623 | 0.042 | 59 | -14.712 | 0.000 | *** |
|  | age | 0.004 | 0.020 | 59 | 0.192 | 0.849 |  |
|  | sexM | 0.032 | 0.117 | 59 | 0.274 | 0.785 |  |
| default | stateSTIv - STIa | -0.269 | 0.133 | 63 | -2.028 | 0.047 | * |
|  | meanFD | 8.996 | 4.447 | 59 | 2.023 | 0.048 | * |
|  | run | -1.004 | 0.078 | 59 | -12.793 | 0.000 | *** |
|  | age | -0.023 | 0.038 | 59 | -0.600 | 0.551 |  |
|  | sexM | 0.114 | 0.217 | 59 | 0.528 | 0.599 |  |
| dorsAtt | stateSTIv - STIa | -0.005 | 0.116 | 63 | -0.043 | 0.966 |  |
|  | meanFD | 4.552 | 2.550 | 59 | 1.785 | 0.079 |  |
|  | run | -0.514 | 0.045 | 59 | -11.426 | 0.000 | *** |
|  | age | -0.017 | 0.022 | 59 | -0.773 | 0.442 |  |
|  | sexM | 0.026 | 0.124 | 59 | 0.213 | 0.832 |  |
| limbic | stateSTIv - STIa | -0.029 | 0.054 | 63 | -0.542 | 0.590 |  |
|  | meanFD | 1.591 | 1.324 | 59 | 1.202 | 0.234 |  |
|  | run | -0.256 | 0.023 | 59 | -10.964 | 0.000 | *** |
|  | age | -0.007 | 0.011 | 59 | -0.637 | 0.527 |  |
|  | sexM | 0.076 | 0.064 | 59 | 1.176 | 0.244 |  |
| somatomotor | stateSTIv - STIa | -0.107 | 0.122 | 63 | -0.882 | 0.381 |  |
|  | meanFD | 5.824 | 3.774 | 59 | 1.543 | 0.128 |  |
|  | run | -0.728 | 0.067 | 59 | -10.932 | 0.000 | *** |
|  | age | -0.029 | 0.032 | 59 | -0.917 | 0.363 |  |
|  | sexM | 0.135 | 0.184 | 59 | 0.733 | 0.466 |  |
| ventrAtt | stateSTIv - STIa | -0.011 | 0.091 | 63 | -0.122 | 0.903 |  |
|  | meanFD | 4.649 | 2.102 | 59 | 2.211 | 0.031 | * |
|  | run | -0.383 | 0.037 | 59 | -10.339 | 0.000 | *** |
|  | age | -0.019 | 0.018 | 59 | -1.080 | 0.285 |  |
|  | sexM | 0.070 | 0.102 | 59 | 0.685 | 0.496 |  |
| visual | stateSTIv - STIa | 0.028 | 0.108 | 63 | 0.260 | 0.796 |  |
|  | meanFD | 3.988 | 2.833 | 59 | 1.408 | 0.164 |  |
|  | run | -0.581 | 0.050 | 59 | -11.623 | 0.000 | *** |
|  | age | 0.010 | 0.024 | 59 | 0.436 | 0.665 |  |
|  | sexM | 0.112 | 0.138 | 59 | 0.814 | 0.419 |  |

lme per within-network combination: euclidean distance = STc vs. STi + confounds. Build-in Matlab function \*pdist\* with the parameter 'euclidean' was used to calculate the Euclidean distance. Task-activity regressed out, schaefer200 parcellation

**Table S12.** Bayes paired t-test between FC similarity (euclidean distance) of the single task modalitiy pairings.

|  | network | Bayes Factor | error |
| --- | --- | --- | --- |
| between network | control | 0.140 | 0.000905 |
|  | default | 0.216 | 0.000680 |
|  | dorsAtt | 0.174 | 0.000784 |
|  | limbic | 0.178 | 0.000773 |
|  | somatomotor | 0.145 | 0.000885 |
|  | ventrAtt | 0.172 | 0.000791 |
|  | visual | 0.153 | 0.000854 |
| within network | control | 0.203 | 0.000709 |
|  | default | 0.988 | 0.000231 |
|  | dorsAtt | 0.137 | 0.000916 |
|  | limbic | 0.158 | 0.000834 |
|  | somatomotor | 0.201 | 0.000713 |
|  | ventrAtt | 0.138 | 0.000912 |
|  | visual | 0.142 | 0.000897 |
| Task-activity regressed out, schaefer200 parcellation |  |  |  |

The model testing within the default-mode network showed as significant effect (*p* = .047, uncorrected) for the factor modality pairing (stateSTIv-STIa), but the corresponding Bayes Factor still providing anecdotal evidence for the null hypothesis (*M* = 0.26, 95% HDI [−0.01, 0.50], BF_10_ = 0.99).

### FC similarity between single and dual tasks

**Table S13.** FC similarity (cosine distance) between single and dual task per between-network combination, full model results.

|  |  | value | std. error | df | t-value | p-value |  |
| --- | --- | --- | --- | --- | --- | --- | --- |
| control | stateSTc-STi | -0.629 | 0.026 | 42 | -23.806 | 0.000 | *** |
|  | meanFD | -0.113 | 0.209 | 42 | -0.543 | 0.590 |  |
|  | run | -0.020 | 0.005 | 42 | -4.349 | 0.000 | *** |
|  | age | -0.001 | 0.002 | 63 | -0.616 | 0.540 |  |
|  | sexM | -0.006 | 0.011 | 63 | -0.519 | 0.605 |  |
| default | stateSTc-STi | -0.625 | 0.028 | 42 | -21.968 | 0.000 | *** |
|  | meanFD | 0.119 | 0.238 | 42 | 0.501 | 0.619 |  |
|  | run | -0.020 | 0.005 | 42 | -3.971 | 0.000 | *** |
|  | age | -0.002 | 0.002 | 63 | -0.826 | 0.412 |  |
|  | sexM | 0.005 | 0.013 | 63 | 0.355 | 0.724 |  |
| dorsAtt | stateSTc-STi | -0.626 | 0.032 | 42 | -19.413 | 0.000 | *** |
|  | meanFD | 0.047 | 0.255 | 42 | 0.185 | 0.854 |  |
|  | run | -0.019 | 0.006 | 42 | -3.328 | 0.002 | ** |
|  | age | -0.002 | 0.002 | 63 | -0.985 | 0.328 |  |
|  | sexM | 0.000 | 0.013 | 63 | 0.015 | 0.988 |  |
| limbic | stateSTc-STi | -0.665 | 0.030 | 42 | -22.343 | 0.000 | *** |
|  | meanFD | 0.057 | 0.238 | 42 | 0.238 | 0.813 |  |
|  | run | -0.017 | 0.005 | 42 | -3.238 | 0.002 | ** |
|  | age | -0.003 | 0.002 | 63 | -1.159 | 0.251 |  |
|  | sexM | 0.005 | 0.012 | 63 | 0.398 | 0.692 |  |
| somatomotor | stateSTc-STi | -0.678 | 0.030 | 42 | -22.387 | 0.000 | *** |
|  | meanFD | 0.070 | 0.239 | 42 | 0.292 | 0.772 |  |
|  | run | -0.015 | 0.005 | 42 | -2.846 | 0.007 | ** |
|  | age | -0.001 | 0.002 | 63 | -0.431 | 0.668 |  |
|  | sexM | 0.004 | 0.012 | 63 | 0.330 | 0.742 |  |
| ventrAtt | stateSTc-STi | -0.666 | 0.035 | 42 | -18.957 | 0.000 | *** |
|  | meanFD | 0.228 | 0.277 | 42 | 0.824 | 0.414 |  |
|  | run | -0.015 | 0.006 | 42 | -2.383 | 0.022 | * |
|  | age | 0.001 | 0.003 | 63 | 0.288 | 0.774 |  |
|  | sexM | -0.001 | 0.014 | 63 | -0.050 | 0.961 |  |
| visual | stateSTc-STi | -0.706 | 0.030 | 42 | -23.777 | 0.000 | *** |
|  | meanFD | 0.071 | 0.234 | 42 | 0.302 | 0.764 |  |
|  | run | -0.015 | 0.005 | 42 | -2.891 | 0.006 | ** |
|  | age | 0.002 | 0.002 | 63 | 0.733 | 0.466 |  |
|  | sexM | -0.018 | 0.012 | 63 | -1.472 | 0.146 |  |

Time per between-network combination: cosine distance = ST vs. DT + confounds. Between connections averaged for each network. Task-activity regressed out, schaefer200 parcellation

**Table S14.**
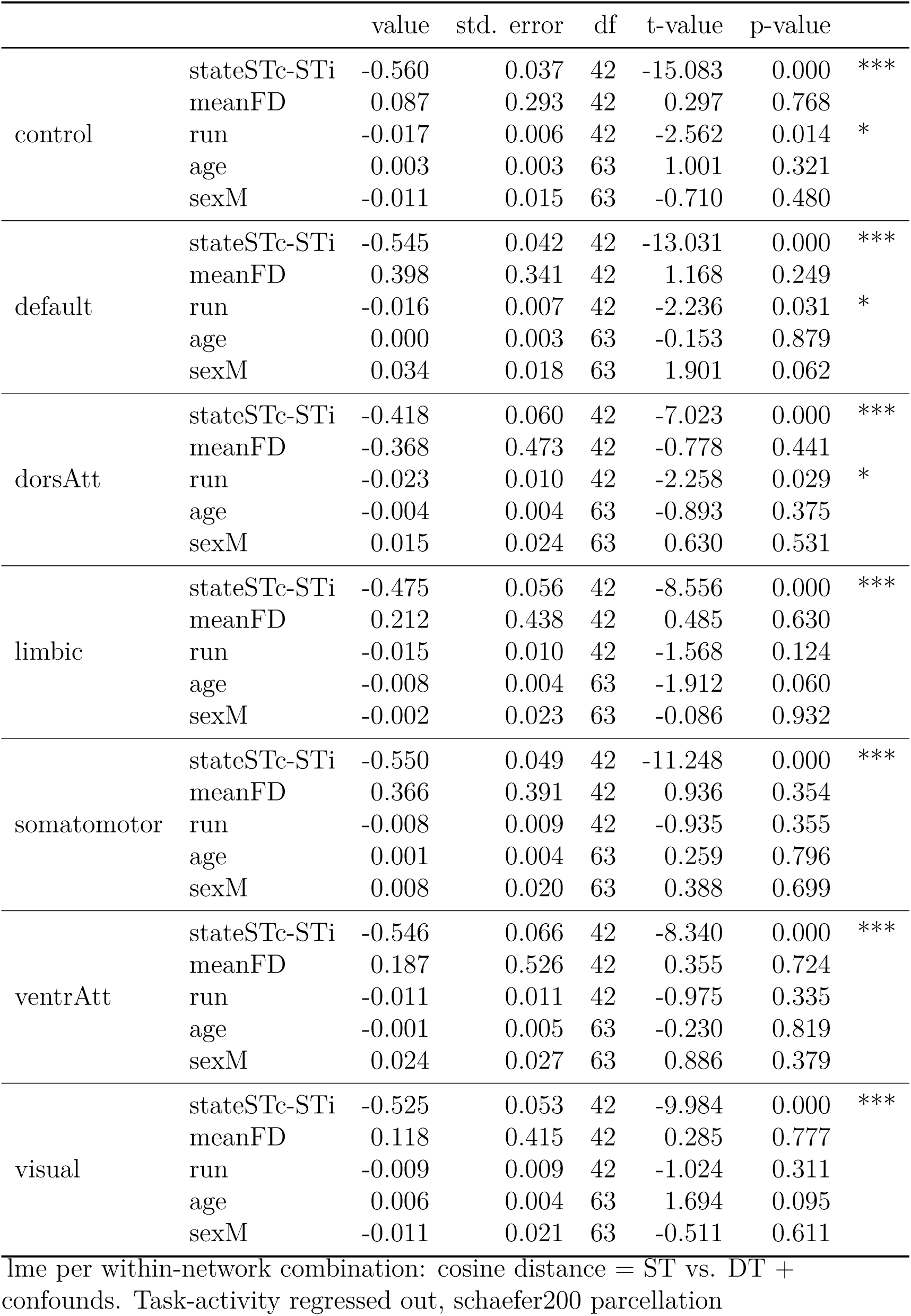
FC similarity (cosine distance) between single and dual task per within-network combination, full model results.

|  |  | value | std. error | df | t-value | p-value |  |
| --- | --- | --- | --- | --- | --- | --- | --- |
| control | stateSTc-STi | -0.560 | 0.037 | 42 | -15.083 | 0.000 | *** |
|  | meanFD | 0.087 | 0.293 | 42 | 0.297 | 0.768 |  |
|  | run | -0.017 | 0.006 | 42 | -2.562 | 0.014 | * |
|  | age | 0.003 | 0.003 | 63 | 1.001 | 0.321 |  |
|  | sexM | -0.011 | 0.015 | 63 | -0.710 | 0.480 |  |
| default | stateSTc-STi | -0.545 | 0.042 | 42 | -13.031 | 0.000 | *** |
|  | meanFD | 0.398 | 0.341 | 42 | 1.168 | 0.249 |  |
|  | run | -0.016 | 0.007 | 42 | -2.236 | 0.031 | * |
|  | age | 0.000 | 0.003 | 63 | -0.153 | 0.879 |  |
|  | sexM | 0.034 | 0.018 | 63 | 1.901 | 0.062 |  |
| dorsAtt | stateSTc-STi | -0.418 | 0.060 | 42 | -7.023 | 0.000 | *** |
|  | meanFD | -0.368 | 0.473 | 42 | -0.778 | 0.441 |  |
|  | run | -0.023 | 0.010 | 42 | -2.258 | 0.029 | * |
|  | age | -0.004 | 0.004 | 63 | -0.893 | 0.375 |  |
|  | sexM | 0.015 | 0.024 | 63 | 0.630 | 0.531 |  |
| limbic | stateSTc-STi | -0.475 | 0.056 | 42 | -8.556 | 0.000 | *** |
|  | meanFD | 0.212 | 0.438 | 42 | 0.485 | 0.630 |  |
|  | run | -0.015 | 0.010 | 42 | -1.568 | 0.124 |  |
|  | age | -0.008 | 0.004 | 63 | -1.912 | 0.060 |  |
|  | sexM | -0.002 | 0.023 | 63 | -0.086 | 0.932 |  |
| somatomotor | stateSTc-STi | -0.550 | 0.049 | 42 | -11.248 | 0.000 | *** |
|  | meanFD | 0.366 | 0.391 | 42 | 0.936 | 0.354 |  |
|  | run | -0.008 | 0.009 | 42 | -0.935 | 0.355 |  |
|  | age | 0.001 | 0.004 | 63 | 0.259 | 0.796 |  |
|  | sexM | 0.008 | 0.020 | 63 | 0.388 | 0.699 |  |
| ventrAtt | stateSTc-STi | -0.546 | 0.066 | 42 | -8.340 | 0.000 | *** |
|  | meanFD | 0.187 | 0.526 | 42 | 0.355 | 0.724 |  |
|  | run | -0.011 | 0.011 | 42 | -0.975 | 0.335 |  |
|  | age | -0.001 | 0.005 | 63 | -0.230 | 0.819 |  |
|  | sexM | 0.024 | 0.027 | 63 | 0.886 | 0.379 |  |
| visual | stateSTc-STi | -0.525 | 0.053 | 42 | -9.984 | 0.000 | *** |
|  | meanFD | 0.118 | 0.415 | 42 | 0.285 | 0.777 |  |
|  | run | -0.009 | 0.009 | 42 | -1.024 | 0.311 |  |
|  | age | 0.006 | 0.004 | 63 | 1.694 | 0.095 |  |
|  | sexM | -0.011 | 0.021 | 63 | -0.511 | 0.611 |  |

line per within-network combination: cosine distance = ST vs. DT + confounds. Task-activity regressed out, schaefer200 parcellation

**Table S15.** Bayes paired t-test between similarity (cosine distance) of single and dual tasks.

|  | network | Bayes Factor | error |
| --- | --- | --- | --- |
| between network | control | 4.02e+38 | 0 |
|  | default | 2.91e+39 | 0 |
|  | dorsAtt | 2.94e+35 | 0 |
|  | limbic | 1.43e+38 | 0 |
|  | somatomotor | 6.34e+37 | 0 |
|  | ventrAtt | 2.19e+34 | 0 |
|  | visual | 8.37e+38 | 0 |
| within network | control | 3.63e+30 | 0 |
|  | default | 3.04e+29 | 0 |
|  | dorsAtt | 3.96e+19 | 0 |
|  | limbic | 5.82e+21 | 0 |
|  | somatomotor | 9.72e+24 | 0 |
|  | ventrAtt | 2.40e+20 | 0 |
|  | visual | 2.03e+22 | 0 |
| Task-activity regressed out, schaefer200 parcellation |  |  |  |

### Functional connectivity between dual tasks

**Table S16.**
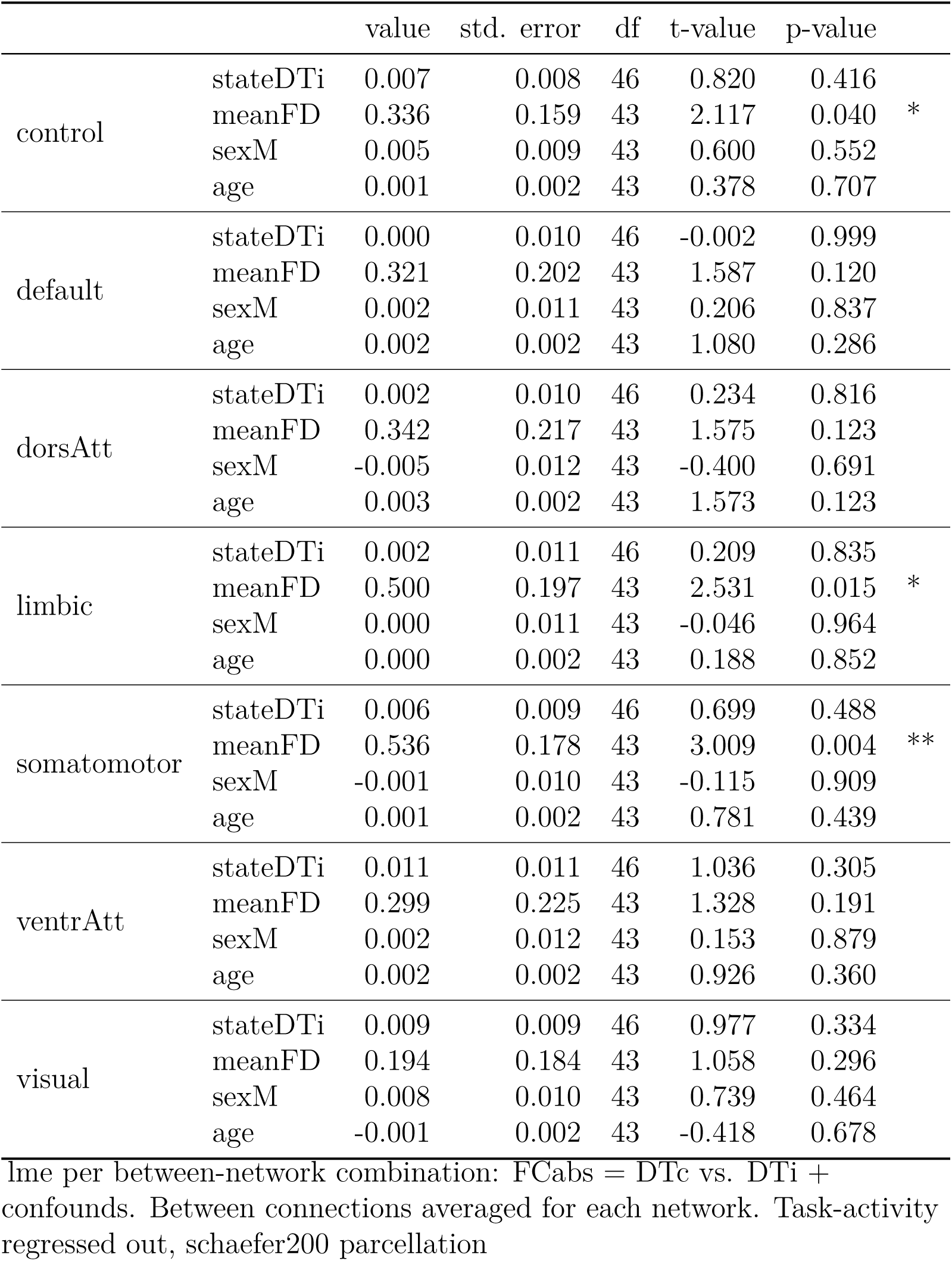
Functional connectivity (absolute) between dual tasks per between-network combination, full model results.

**Table S17.** Functional connectivity (absolute) between dual tasks per within-network combination, full model results.

|  |  | value | std. error | df | t-value | p-value |
| --- | --- | --- | --- | --- | --- | --- |
| control | stateDTi | 0.009 | 0.012 | 46 | 0.759 | 0.452 |
|  | meanFD | 0.180 | 0.229 | 43 | 0.784 | 0.437 |
|  | sexM | 0.007 | 0.013 | 43 | 0.583 | 0.563 |
|  | age | 0.001 | 0.002 | 43 | 0.346 | 0.731 |
| default | stateDTi | -0.019 | 0.014 | 46 | -1.370 | 0.177 |
|  | meanFD | 0.178 | 0.327 | 43 | 0.546 | 0.588 |
|  | sexM | -0.008 | 0.018 | 43 | -0.436 | 0.665 |
|  | age | 0.002 | 0.003 | 43 | 0.769 | 0.446 |
| dorsAtt | stateDTi | 0.006 | 0.022 | 46 | 0.290 | 0.773 |
|  | meanFD | 0.418 | 0.513 | 43 | 0.815 | 0.419 |
|  | sexM | -0.029 | 0.028 | 43 | -1.031 | 0.308 |
|  | age | 0.006 | 0.005 | 43 | 1.181 | 0.244 |
| limbic | stateDTi | -0.002 | 0.023 | 46 | -0.093 | 0.926 |
|  | meanFD | 0.521 | 0.423 | 43 | 1.231 | 0.225 |
|  | sexM | 0.011 | 0.023 | 43 | 0.465 | 0.644 |
|  | age | -0.003 | 0.004 | 43 | -0.794 | 0.431 |
| somatomotor | stateDTi | -0.008 | 0.019 | 46 | -0.425 | 0.673 |
|  | meanFD | 0.308 | 0.350 | 43 | 0.878 | 0.385 |
|  | sexM | -0.031 | 0.019 | 43 | -1.622 | 0.112 |
|  | age | 0.002 | 0.003 | 43 | 0.536 | 0.595 |
| ventrAtt | stateDTi | 0.036 | 0.025 | 46 | 1.465 | 0.150 |
|  | meanFD | 0.146 | 0.528 | 43 | 0.277 | 0.783 |
|  | sexM | -0.018 | 0.029 | 43 | -0.607 | 0.547 |
|  | age | 0.005 | 0.005 | 43 | 0.961 | 0.342 |
| visual | stateDTi | 0.027 | 0.025 | 46 | 1.078 | 0.287 |
|  | meanFD | 0.125 | 0.500 | 43 | 0.250 | 0.804 |
|  | sexM | 0.020 | 0.028 | 43 | 0.719 | 0.476 |
|  | age | -0.005 | 0.005 | 43 | -1.091 | 0.281 |

lme per within-network combination: FCabs = DTc vs. DTi + confounds. Task-activity regressed out, schaefer200 parcellation

**Table S18.** Bayes paired t-test between absolute functional connectivity of dual task modalitiy pairings.

|  | network | Bayes Factor | error |
| --- | --- | --- | --- |
| between network | control | 0.220 | 0.000546 |
|  | default | 0.158 | 0.000638 |
|  | dorsAtt | 0.163 | 0.000630 |
|  | limbic | 0.162 | 0.000632 |
|  | somatomotor | 0.201 | 0.000570 |
|  | ventrAtt | 0.267 | 0.000495 |
|  | visual | 0.252 | 0.000510 |
| within network | control | 0.210 | 0.000559 |
|  | default | 0.390 | 0.000404 |
|  | dorsAtt | 0.165 | 0.000626 |
|  | limbic | 0.159 | 0.000637 |
|  | somatomotor | 0.173 | 0.000613 |
|  | ventrAtt | 0.443 | 0.000377 |
|  | visual | 0.278 | 0.000485 |
| Task-activity regressed out, schaefer200 parcellation |  |  |  |

### Functional connectivity: relative Values

In addition to the absolute connectivity values, which neglect the information if regions are positively or negatively correlated, we repeated the analysis with the relative connectivity values.

**Table S19.** Functional connectivity (relative) between dual tasks per between-network combination, full model results.

|  |  | value | std. error | df | t-value | p-value |
| --- | --- | --- | --- | --- | --- | --- |
| control | stateDTi | -0.005 | 0.008 | 46 | -0.667 | 0.508 |
|  | meanFD | 0.108 | 0.211 | 43 | 0.510 | 0.612 |
|  | sexM | 0.008 | 0.012 | 43 | 0.659 | 0.513 |
|  | age | 0.002 | 0.002 | 43 | 0.782 | 0.439 |
| default | stateDTi | 0.011 | 0.010 | 46 | 1.107 | 0.274 |
|  | meanFD | 0.194 | 0.238 | 43 | 0.812 | 0.421 |
|  | sexM | 0.011 | 0.013 | 43 | 0.812 | 0.421 |
|  | age | 0.001 | 0.002 | 43 | 0.501 | 0.619 |
| dorsAtt | stateDTi | -0.002 | 0.010 | 46 | -0.188 | 0.852 |
|  | meanFD | 0.120 | 0.219 | 43 | 0.547 | 0.587 |
|  | sexM | 0.010 | 0.012 | 43 | 0.834 | 0.409 |
|  | age | 0.001 | 0.002 | 43 | 0.345 | 0.732 |
| limbic | stateDTi | -0.005 | 0.013 | 46 | -0.357 | 0.722 |
|  | meanFD | -0.344 | 0.242 | 43 | -1.422 | 0.162 |
|  | sexM | 0.013 | 0.013 | 43 | 0.962 | 0.341 |
|  | age | -0.001 | 0.002 | 43 | -0.270 | 0.788 |
| somatomotor | stateDTi | -0.004 | 0.009 | 46 | -0.387 | 0.701 |
|  | meanFD | 0.033 | 0.191 | 43 | 0.171 | 0.865 |
|  | sexM | -0.002 | 0.011 | 43 | -0.181 | 0.857 |
|  | age | 0.002 | 0.002 | 43 | 1.096 | 0.279 |
| ventrAtt | stateDTi | 0.009 | 0.010 | 46 | 0.943 | 0.351 |
|  | meanFD | -0.039 | 0.203 | 43 | -0.190 | 0.850 |
|  | sexM | 0.002 | 0.011 | 43 | 0.205 | 0.839 |
|  | age | 0.000 | 0.002 | 43 | -0.033 | 0.974 |
| visual | stateDTi | -0.011 | 0.011 | 46 | -0.963 | 0.341 |
|  | meanFD | 0.209 | 0.238 | 43 | 0.878 | 0.385 |
|  | sexM | 0.001 | 0.013 | 43 | 0.040 | 0.968 |
|  | age | 0.000 | 0.002 | 43 | 0.097 | 0.923 |

lme per between-network combination: FCrel = DTc vs. DTi + confounds. Between connections averaged for each network. Task-activity regressed out, schaefer200 parcellation

**Table S20.** Functional connectivity (relative) between dual tasks per within-network combination, full model results.

|  |  | value | std. error | df | t-value | p-value |
| --- | --- | --- | --- | --- | --- | --- |
| control | stateDTi | 0.011 | 0.020 | 46 | 0.553 | 0.583 |
|  | meanFD | -0.146 | 0.396 | 43 | -0.370 | 0.713 |
|  | sexM | 0.024 | 0.022 | 43 | 1.104 | 0.276 |
|  | age | -0.007 | 0.004 | 43 | -1.745 | 0.088 |
| default | stateDTi | -0.024 | 0.021 | 46 | -1.156 | 0.254 |
|  | meanFD | -0.515 | 0.494 | 43 | -1.042 | 0.303 |
|  | sexM | -0.050 | 0.027 | 43 | -1.815 | 0.077 |
|  | age | 0.004 | 0.005 | 43 | 0.821 | 0.416 |
| dorsAtt | stateDTi | 0.018 | 0.035 | 46 | 0.505 | 0.616 |
|  | meanFD | 0.742 | 0.838 | 43 | 0.886 | 0.381 |
|  | sexM | -0.039 | 0.046 | 43 | -0.849 | 0.401 |
|  | age | 0.009 | 0.008 | 43 | 1.149 | 0.257 |
| limbic | stateDTi | 0.012 | 0.037 | 46 | 0.328 | 0.744 |
|  | meanFD | -0.072 | 0.678 | 43 | -0.106 | 0.916 |
|  | sexM | -0.056 | 0.038 | 43 | -1.480 | 0.146 |
|  | age | 0.007 | 0.007 | 43 | 1.129 | 0.265 |
| somatomotor | stateDTi | -0.008 | 0.033 | 46 | -0.249 | 0.805 |
|  | meanFD | 0.055 | 0.691 | 43 | 0.079 | 0.937 |
|  | sexM | -0.049 | 0.038 | 43 | -1.289 | 0.204 |
|  | age | -0.007 | 0.007 | 43 | -0.992 | 0.327 |
| ventrAtt | stateDTi | 0.031 | 0.039 | 46 | 0.807 | 0.424 |
|  | meanFD | -0.220 | 0.899 | 43 | -0.244 | 0.808 |
|  | sexM | -0.058 | 0.050 | 43 | -1.166 | 0.250 |
|  | age | 0.000 | 0.009 | 43 | -0.021 | 0.983 |
| visual | stateDTi | 0.043 | 0.036 | 46 | 1.207 | 0.234 |
|  | meanFD | -0.463 | 0.784 | 43 | -0.590 | 0.558 |
|  | sexM | 0.055 | 0.043 | 43 | 1.278 | 0.208 |
|  | age | -0.014 | 0.008 | 43 | -1.850 | 0.071 |

lme per within-network combination: FCrel = DTc vs. DTi + confounds. Task-activity regressed out, schaefer200 parcellation

**Table S21.** Bayes paired t-test between relative functional connectivity of dual task modalitiy pairings.

|  | network | Bayes Factor | error |
| --- | --- | --- | --- |
| between network | control | 0.197 | 0.000576 |
|  | default | 0.287 | 0.000477 |
|  | dorsAtt | 0.161 | 0.000633 |
|  | limbic | 0.168 | 0.000621 |
|  | somatomotor | 0.170 | 0.000617 |
|  | ventrAtt | 0.244 | 0.000518 |
|  | visual | 0.248 | 0.000513 |
| within network | control | 0.184 | 0.000595 |
|  | default | 0.302 | 0.000464 |
|  | dorsAtt | 0.179 | 0.000602 |
|  | limbic | 0.166 | 0.000624 |
|  | somatomotor | 0.163 | 0.000629 |
|  | ventrAtt | 0.217 | 0.000549 |
|  | visual | 0.320 | 0.000450 |
| Task-activity regressed out, schaefer200 parcellation |  |  |  |

### Brain-Behavior correlation

**Figure S2.**
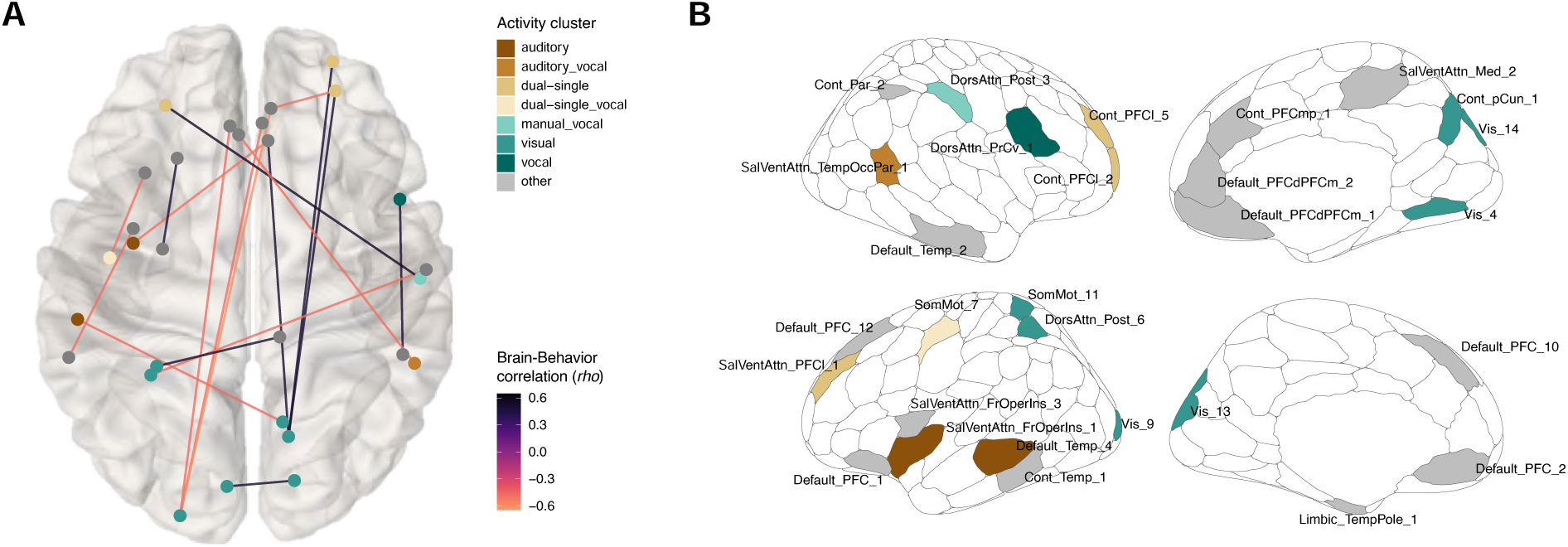
Relation between dual-task behavior and functional connectivity. **A**: The superior view of the brain with the connections which are significantly related to the dual-task behavior (correlated with p < .001 and without the restriction of the region overlapping with task-relevant cluster). Color of regions indicate the corresponding activity cluster, color of connections the strengh of the correlation. The rho value is based on the partial correlation, corrected for age, gender and framewise displacement. **B**: Anatomical location of relevant schaefer200 regions for brain-behavior relation, together with the task-based activity clusters. Labels correspond to the Schaefer regions (name of the region according to schaefer200 7networks), colors to the activity clusters, while grey indicate no overlap between Schaefer region and our task-based activity clusters.

### Preprocessing: schaefer100 parcellation

We additionally ruled out that the results are specific to one parcellation, by repeating the main analysis with 100, instead of 200 regions, defined by Schaefer et al. (2018). All other preprocessing steps were the same as reported in the manuscript. We first report the FC similarity between modality pairings in single tasks and then the FC strength between dual tasks.

**Table S22.** FC similarity (cosine distance) between single tasks per between-network combination, full model results.

|  |  | value | std. error | df | t-value | p-value |  |
| --- | --- | --- | --- | --- | --- | --- | --- |
| control | stateSTIv - STIa | -0.001 | 0.007 | 63 | -0.183 | 0.855 |  |
|  | meanFD | 0.266 | 0.239 | 59 | 1.114 | 0.270 |  |
|  | run | -0.021 | 0.004 | 59 | -4.946 | 0.000 | *** |
|  | age | -0.001 | 0.002 | 59 | -0.569 | 0.571 |  |
|  | sexM | -0.017 | 0.012 | 59 | -1.456 | 0.151 |  |
| default | stateSTIv - STIa | 0.002 | 0.005 | 63 | 0.367 | 0.715 |  |
|  | meanFD | 0.437 | 0.235 | 59 | 1.863 | 0.067 |  |
|  | run | -0.016 | 0.004 | 59 | -3.980 | 0.000 | *** |
|  | age | -0.001 | 0.002 | 59 | -0.392 | 0.697 |  |
|  | sexM | -0.005 | 0.011 | 59 | -0.470 | 0.640 |  |
| dorsAtt | stateSTIv - STIa | -0.007 | 0.007 | 63 | -1.103 | 0.274 |  |
|  | meanFD | 0.527 | 0.237 | 59 | 2.221 | 0.030 | * |
|  | run | -0.017 | 0.004 | 59 | -4.097 | 0.000 | *** |
|  | age | -0.001 | 0.002 | 59 | -0.426 | 0.672 |  |
|  | sexM | -0.016 | 0.012 | 59 | -1.422 | 0.160 |  |
| limbic | stateSTIv - STIa | -0.018 | 0.010 | 63 | -1.751 | 0.085 |  |
|  | meanFD | 0.826 | 0.417 | 59 | 1.979 | 0.052 |  |
|  | run | -0.012 | 0.007 | 59 | -1.567 | 0.122 |  |
|  | age | -0.003 | 0.004 | 59 | -0.971 | 0.335 |  |
|  | sexM | -0.002 | 0.020 | 59 | -0.103 | 0.918 |  |
| somatomotor | stateSTIv - STIa | -0.002 | 0.007 | 63 | -0.212 | 0.833 |  |
|  | meanFD | 0.327 | 0.295 | 59 | 1.107 | 0.273 |  |
|  | run | -0.016 | 0.005 | 59 | -2.981 | 0.004 | ** |
|  | age | -0.005 | 0.003 | 59 | -1.800 | 0.077 |  |
|  | sexM | -0.012 | 0.014 | 59 | -0.858 | 0.395 |  |
| ventrAtt | stateSTIv - STIa | -0.005 | 0.006 | 63 | -0.894 | 0.375 |  |
|  | meanFD | 0.217 | 0.236 | 59 | 0.919 | 0.362 |  |
|  | run | -0.019 | 0.004 | 59 | -4.515 | 0.000 | *** |
|  | age | -0.002 | 0.002 | 59 | -0.764 | 0.448 |  |
|  | sexM | -0.006 | 0.012 | 59 | -0.509 | 0.613 |  |
| visual | stateSTIv - STIa | -0.013 | 0.008 | 63 | -1.681 | 0.098 |  |
|  | meanFD | 0.035 | 0.296 | 59 | 0.120 | 0.905 |  |
|  | run | -0.016 | 0.005 | 59 | -3.072 | 0.003 | ** |
|  | age | -0.004 | 0.003 | 59 | -1.664 | 0.101 |  |
|  | sexM | -0.013 | 0.014 | 59 | -0.896 | 0.374 |  |

lme per between-network combination: cosine distance = STc vs. STi + confounds. Between connections averaged for each network. Task-activity remained in data, schaefer100 parcellation

**Table S23.** FC similarity (cosine distance) between single tasks per within-network combination, full model results.

|  |  | value | std. error | df | t-value | p-value |  |
| --- | --- | --- | --- | --- | --- | --- | --- |
| control | stateSTIv - STIa | 0.009 | 0.006 | 63 | 1.442 | 0.154 |  |
|  | meanFD | 0.070 | 0.213 | 59 | 0.330 | 0.742 |  |
|  | run | -0.018 | 0.004 | 59 | -4.804 | 0.000 | *** |
|  | age | 0.001 | 0.002 | 59 | 0.423 | 0.674 |  |
|  | sexM | -0.006 | 0.010 | 59 | -0.602 | 0.550 |  |
| default | stateSTIv - STIa | 0.000 | 0.003 | 63 | 0.074 | 0.942 |  |
|  | meanFD | 0.359 | 0.185 | 59 | 1.943 | 0.057 |  |
|  | run | -0.019 | 0.003 | 59 | -5.840 | 0.000 | *** |
|  | age | 0.000 | 0.002 | 59 | 0.039 | 0.969 |  |
|  | sexM | 0.002 | 0.009 | 59 | 0.258 | 0.798 |  |
| dorsAtt | stateSTIv - STIa | 0.002 | 0.006 | 63 | 0.313 | 0.756 |  |
|  | meanFD | 0.225 | 0.217 | 59 | 1.034 | 0.306 |  |
|  | run | -0.022 | 0.004 | 59 | -5.712 | 0.000 | *** |
|  | age | 0.000 | 0.002 | 59 | -0.151 | 0.881 |  |
|  | sexM | -0.003 | 0.011 | 59 | -0.247 | 0.806 |  |
| limbic | stateSTIv - STIa | -0.002 | 0.009 | 63 | -0.199 | 0.843 |  |
|  | meanFD | 0.220 | 0.263 | 59 | 0.837 | 0.406 |  |
|  | run | -0.015 | 0.005 | 59 | -3.226 | 0.002 | ** |
|  | age | -0.001 | 0.002 | 59 | -0.345 | 0.732 |  |
|  | sexM | 0.017 | 0.013 | 59 | 1.295 | 0.200 |  |
| somatomotor | stateSTIv - STIa | 0.004 | 0.004 | 63 | 1.043 | 0.301 |  |
|  | meanFD | 0.236 | 0.140 | 59 | 1.682 | 0.098 |  |
|  | run | -0.011 | 0.002 | 59 | -4.367 | 0.000 | *** |
|  | age | -0.002 | 0.001 | 59 | -1.277 | 0.207 |  |
|  | sexM | 0.002 | 0.007 | 59 | 0.251 | 0.803 |  |
| ventrAtt | stateSTIv - STIa | 0.005 | 0.005 | 63 | 1.095 | 0.278 |  |
|  | meanFD | 0.178 | 0.212 | 59 | 0.840 | 0.404 |  |
|  | run | -0.014 | 0.004 | 59 | -3.866 | 0.000 | *** |
|  | age | -0.001 | 0.002 | 59 | -0.351 | 0.727 |  |
|  | sexM | 0.013 | 0.010 | 59 | 1.233 | 0.222 |  |
| visual | stateSTIv - STIa | 0.000 | 0.003 | 63 | 0.034 | 0.973 |  |
|  | meanFD | 0.084 | 0.104 | 59 | 0.804 | 0.425 |  |
|  | run | -0.012 | 0.002 | 59 | -6.603 | 0.000 | *** |
|  | age | 0.001 | 0.001 | 59 | 0.574 | 0.568 |  |
|  | sexM | 0.002 | 0.005 | 59 | 0.428 | 0.670 |  |

lme per within-network combination: cosine distance = STc vs. STi + confounds.
Task-activity remained in data, schaefer100 parcellation

**Table S24.** Bayes paired t-test between FC similarity (cosine distance) of the single task modalitiy pairings.

|  | network | Bayes Factor | error |
| --- | --- | --- | --- |
| between network | control | 0.139 | 0.000907 |
|  | default | 0.146 | 0.000878 |
|  | dorsAtt | 0.249 | 0.000618 |
|  | limbic | 0.604 | 0.000332 |
|  | somatomotor | 0.140 | 0.000904 |
|  | ventrAtt | 0.203 | 0.000709 |
|  | visual | 0.539 | 0.000360 |
| within network | control | 0.378 | 0.000464 |
|  | default | 0.137 | 0.000915 |
|  | dorsAtt | 0.144 | 0.000889 |
|  | limbic | 0.140 | 0.000905 |
|  | somatomotor | 0.234 | 0.000645 |
|  | ventrAtt | 0.247 | 0.000621 |
|  | visual | 0.137 | 0.000916 |
| Task-activity remained in data, schaefer100 parcellation |  |  |  |

**Table S25.** Functional connectivity (absolute) between dual tasks per between-network combination, full model results.

|  |  | value | std. error | df | t-value | p-value |  |
| --- | --- | --- | --- | --- | --- | --- | --- |
| control | stateDTi | 0.002 | 0.009 | 46 | 0.217 | 0.829 |  |
|  | meanFD | 0.295 | 0.170 | 43 | 1.729 | 0.091 |  |
|  | sexM | 0.002 | 0.009 | 43 | 0.197 | 0.845 |  |
|  | age | 0.002 | 0.002 | 43 | 0.967 | 0.339 |  |
| default | stateDTi | -0.003 | 0.011 | 46 | -0.285 | 0.777 |  |
|  | meanFD | 0.287 | 0.216 | 43 | 1.329 | 0.191 |  |
|  | sexM | 0.001 | 0.012 | 43 | 0.063 | 0.950 |  |
|  | age | 0.002 | 0.002 | 43 | 1.024 | 0.312 |  |
| dorsAtt | stateDTi | 0.008 | 0.011 | 46 | 0.676 | 0.503 |  |
|  | meanFD | 0.410 | 0.207 | 43 | 1.980 | 0.054 |  |
|  | sexM | -0.002 | 0.011 | 43 | -0.210 | 0.835 |  |
|  | age | 0.004 | 0.002 | 43 | 1.924 | 0.061 |  |
| limbic | stateDTi | 0.003 | 0.014 | 46 | 0.220 | 0.827 |  |
|  | meanFD | 0.555 | 0.262 | 43 | 2.121 | 0.040 | * |
|  | sexM | -0.014 | 0.014 | 43 | -0.977 | 0.334 |  |
|  | age | 0.001 | 0.003 | 43 | 0.363 | 0.719 |  |
| somatomotor | stateDTi | 0.006 | 0.011 | 46 | 0.549 | 0.585 |  |
|  | meanFD | 0.638 | 0.212 | 43 | 3.012 | 0.004 | ** |
|  | sexM | 0.003 | 0.012 | 43 | 0.244 | 0.808 |  |
|  | age | 0.001 | 0.002 | 43 | 0.374 | 0.710 |  |
| ventrAtt | stateDTi | 0.008 | 0.012 | 46 | 0.637 | 0.527 |  |
|  | meanFD | 0.244 | 0.225 | 43 | 1.088 | 0.283 |  |
|  | sexM | -0.001 | 0.012 | 43 | -0.096 | 0.924 |  |
|  | age | 0.002 | 0.002 | 43 | 0.943 | 0.351 |  |
| visual | stateDTi | 0.003 | 0.010 | 46 | 0.285 | 0.777 |  |
|  | meanFD | 0.201 | 0.202 | 43 | 0.994 | 0.326 |  |
|  | sexM | 0.009 | 0.011 | 43 | 0.768 | 0.447 |  |
|  | age | -0.001 | 0.002 | 43 | -0.715 | 0.479 |  |

lme per between-network combination: FCabs = DTc vs. DTi + confounds. Between connections averaged for each network. Task-activity remained in data, schaefer100 parcellation

**Table S26.** Functional connectivity (absolute) between dual tasks per within-network combination, full model results.

|  |  | value | std. error | df | t-value | p-value |
| --- | --- | --- | --- | --- | --- | --- |
| control | stateDTi | 0.025 | 0.020 | 46 | 1.264 | 0.213 |
|  | meanFD | 0.035 | 0.374 | 43 | 0.092 | 0.927 |
|  | sexM | -0.007 | 0.021 | 43 | -0.322 | 0.749 |
|  | age | 0.002 | 0.004 | 43 | 0.528 | 0.600 |
| default | stateDTi | -0.014 | 0.015 | 46 | -0.953 | 0.346 |
|  | meanFD | 0.104 | 0.365 | 43 | 0.284 | 0.778 |
|  | sexM | -0.006 | 0.020 | 43 | -0.303 | 0.763 |
|  | age | 0.004 | 0.004 | 43 | 0.999 | 0.323 |
| dorsAtt | stateDTi | 0.013 | 0.026 | 46 | 0.478 | 0.635 |
|  | meanFD | 0.795 | 0.537 | 43 | 1.481 | 0.146 |
|  | sexM | -0.031 | 0.030 | 43 | -1.044 | 0.302 |
|  | age | 0.007 | 0.005 | 43 | 1.428 | 0.160 |
| limbic | stateDTi | -0.012 | 0.038 | 46 | -0.316 | 0.753 |
|  | meanFD | 0.961 | 0.710 | 43 | 1.354 | 0.183 |
|  | sexM | -0.042 | 0.039 | 43 | -1.063 | 0.294 |
|  | age | 0.002 | 0.007 | 43 | 0.244 | 0.808 |
| somatomotor | stateDTi | -0.013 | 0.026 | 46 | -0.510 | 0.613 |
|  | meanFD | 0.291 | 0.545 | 43 | 0.534 | 0.596 |
|  | sexM | -0.052 | 0.030 | 43 | -1.724 | 0.092 |
|  | age | -0.001 | 0.005 | 43 | -0.207 | 0.837 |
| ventrAtt | stateDTi | 0.037 | 0.031 | 46 | 1.190 | 0.240 |
|  | meanFD | 0.245 | 0.609 | 43 | 0.402 | 0.690 |
|  | sexM | -0.022 | 0.034 | 43 | -0.645 | 0.522 |
|  | age | 0.003 | 0.006 | 43 | 0.529 | 0.600 |
| visual | stateDTi | 0.037 | 0.028 | 46 | 1.324 | 0.192 |
|  | meanFD | -0.139 | 0.599 | 43 | -0.231 | 0.818 |
|  | sexM | 0.020 | 0.033 | 43 | 0.602 | 0.550 |
|  | age | -0.006 | 0.006 | 43 | -1.060 | 0.295 |

lme per within-network combination: FCabs = DTc vs. DTi + confounds. Task-activity remained in data, schaefer100 parcellation

**Table S27.** Bayes paired t-test between absolute functional connectivity of dual task modalitiy pairings.

|  | network | Bayes Factor | error |
| --- | --- | --- | --- |
| between network | control | 0.162 | 0.000632 |
|  | default | 0.165 | 0.000626 |
|  | dorsAtt | 0.196 | 0.000577 |
|  | limbic | 0.161 | 0.000633 |
|  | somatomotor | 0.183 | 0.000596 |
|  | ventrAtt | 0.193 | 0.000581 |
|  | visual | 0.165 | 0.000626 |
| within network | control | 0.333 | 0.000441 |
|  | default | 0.246 | 0.000516 |
|  | dorsAtt | 0.177 | 0.000606 |
|  | limbic | 0.165 | 0.000626 |
|  | somatomotor | 0.180 | 0.000601 |
|  | ventrAtt | 0.314 | 0.000455 |
|  | visual | 0.368 | 0.000418 |
| Task-activity remained in data, schaefer100 parcellation |  |  |  |

### Preprocessing: neural-task activity remained

To test the robustness of our findings, we ruled out that the non-significant results are due to the regression of task-related activity in the preprocessing step. Accordingly, we repeated the pre-registered analysis with timeseries data where neural-task activity is still remaining. All the other preprocessing steps were the same as reported in the manuscript. We first report the FC similarity between modality pairings in single tasks and then the FC strength between dual tasks.

**Table S28.** FC similarity (cosine distance) between single tasks per between-network combination, full model results.

|  |  | value | std. error | df | t-value | p-value |  |
| --- | --- | --- | --- | --- | --- | --- | --- |
| control | stateSTIv - STIa | 0.005 | 0.008 | 63 | 0.644 | 0.522 |  |
|  | meanFD | 0.354 | 0.242 | 59 | 1.460 | 0.150 |  |
|  | run | -0.030 | 0.004 | 59 | -6.948 | 0.000 | *** |
|  | age | -0.001 | 0.002 | 59 | -0.564 | 0.575 |  |
|  | sexM | 0.015 | 0.012 | 59 | 1.252 | 0.216 |  |
| default | stateSTIv - STIa | 0.002 | 0.008 | 63 | 0.194 | 0.846 |  |
|  | meanFD | 0.472 | 0.246 | 59 | 1.921 | 0.060 |  |
|  | run | -0.026 | 0.004 | 59 | -6.038 | 0.000 | *** |
|  | age | -0.001 | 0.002 | 59 | -0.517 | 0.607 |  |
|  | sexM | 0.016 | 0.012 | 59 | 1.316 | 0.193 |  |
| dorsAtt | stateSTIv - STIa | 0.012 | 0.008 | 63 | 1.489 | 0.142 |  |
|  | meanFD | 0.456 | 0.234 | 59 | 1.949 | 0.056 |  |
|  | run | -0.026 | 0.004 | 59 | -6.352 | 0.000 | *** |
|  | age | -0.001 | 0.002 | 59 | -0.461 | 0.646 |  |
|  | sexM | 0.018 | 0.011 | 59 | 1.595 | 0.116 |  |
| limbic | stateSTIv - STIa | 0.000 | 0.011 | 63 | -0.032 | 0.974 |  |
|  | meanFD | 0.439 | 0.363 | 59 | 1.209 | 0.232 |  |
|  | run | -0.025 | 0.006 | 59 | -3.928 | 0.000 | *** |
|  | age | -0.001 | 0.003 | 59 | -0.478 | 0.635 |  |
|  | sexM | 0.044 | 0.018 | 59 | 2.461 | 0.017 | * |
| somatomotor | stateSTIv - STIa | 0.008 | 0.010 | 63 | 0.795 | 0.430 |  |
|  | meanFD | 0.132 | 0.317 | 59 | 0.418 | 0.678 |  |
|  | run | -0.029 | 0.006 | 59 | -5.170 | 0.000 | *** |
|  | age | -0.002 | 0.003 | 59 | -0.753 | 0.455 |  |
|  | sexM | -0.004 | 0.015 | 59 | -0.232 | 0.817 |  |
| ventrAtt | stateSTIv - STIa | -0.002 | 0.010 | 63 | -0.192 | 0.849 |  |
|  | meanFD | 0.532 | 0.233 | 59 | 2.278 | 0.026 | * |
|  | run | -0.028 | 0.004 | 59 | -6.744 | 0.000 | *** |
|  | age | -0.001 | 0.002 | 59 | -0.430 | 0.669 |  |
|  | sexM | 0.010 | 0.011 | 59 | 0.864 | 0.391 |  |
| visual | stateSTIv - STIa | -0.002 | 0.012 | 63 | -0.148 | 0.883 |  |
|  | meanFD | 0.481 | 0.332 | 59 | 1.448 | 0.153 |  |
|  | run | -0.030 | 0.006 | 59 | -5.114 | 0.000 | *** |
|  | age | 0.000 | 0.003 | 59 | -0.087 | 0.931 |  |
|  | sexM | 0.031 | 0.016 | 59 | 1.937 | 0.058 |  |

lme per between-network combination: cosine distance = STc vs. STi + confounds. Between connections averaged for each network. Task-activity remained in data, schaefer200 parcellation

**Table S29.** FC similarity (cosine distance) between single tasks per within-network combination, full model results.

|  |  | value | std. error | df | t-value | p-value |  |
| --- | --- | --- | --- | --- | --- | --- | --- |
| control | stateSTIv - STIa | 0.009 | 0.005 | 63 | 1.676 | 0.099 |  |
|  | meanFD | 0.211 | 0.209 | 59 | 1.011 | 0.316 |  |
|  | run | -0.029 | 0.004 | 59 | -7.944 | 0.000 | *** |
|  | age | 0.001 | 0.002 | 59 | 0.745 | 0.459 |  |
|  | sexM | 0.001 | 0.010 | 59 | 0.117 | 0.907 |  |
| default | stateSTIv - STIa | 0.005 | 0.006 | 63 | 0.793 | 0.431 |  |
|  | meanFD | 0.319 | 0.202 | 59 | 1.582 | 0.119 |  |
|  | run | -0.022 | 0.004 | 59 | -6.136 | 0.000 | *** |
|  | age | -0.001 | 0.002 | 59 | -0.688 | 0.494 |  |
|  | sexM | 0.019 | 0.010 | 59 | 1.938 | 0.057 |  |
| dorsAtt | stateSTIv - STIa | 0.013 | 0.005 | 63 | 2.767 | 0.007 | ** |
|  | meanFD | 0.187 | 0.187 | 59 | 0.999 | 0.322 |  |
|  | run | -0.015 | 0.003 | 59 | -4.507 | 0.000 | *** |
|  | age | 0.000 | 0.002 | 59 | 0.176 | 0.861 |  |
|  | sexM | 0.008 | 0.009 | 59 | 0.845 | 0.402 |  |
| limbic | stateSTIv - STIa | 0.000 | 0.006 | 63 | 0.075 | 0.941 |  |
|  | meanFD | 0.064 | 0.184 | 59 | 0.350 | 0.728 |  |
|  | run | -0.014 | 0.003 | 59 | -4.343 | 0.000 | *** |
|  | age | 0.000 | 0.002 | 59 | 0.110 | 0.912 |  |
|  | sexM | 0.018 | 0.009 | 59 | 2.057 | 0.044 | * |
| somatomotor | stateSTIv - STIa | 0.002 | 0.005 | 63 | 0.370 | 0.713 |  |
|  | meanFD | 0.101 | 0.199 | 59 | 0.509 | 0.613 |  |
|  | run | -0.018 | 0.004 | 59 | -5.235 | 0.000 | *** |
|  | age | 0.001 | 0.002 | 59 | 0.423 | 0.674 |  |
|  | sexM | -0.007 | 0.010 | 59 | -0.768 | 0.445 |  |
| ventrAtt | stateSTIv - STIa | 0.001 | 0.004 | 63 | 0.183 | 0.856 |  |
|  | meanFD | 0.079 | 0.158 | 59 | 0.503 | 0.617 |  |
|  | run | -0.018 | 0.003 | 59 | -6.333 | 0.000 | *** |
|  | age | -0.001 | 0.001 | 59 | -0.490 | 0.626 |  |
|  | sexM | -0.001 | 0.008 | 59 | -0.095 | 0.924 |  |
| visual | stateSTIv - STIa | 0.000 | 0.003 | 63 | 0.084 | 0.933 |  |
|  | meanFD | 0.030 | 0.137 | 59 | 0.216 | 0.829 |  |
|  | run | -0.017 | 0.002 | 59 | -6.992 | 0.000 | *** |
|  | age | 0.000 | 0.001 | 59 | 0.422 | 0.675 |  |
|  | sexM | 0.001 | 0.007 | 59 | 0.163 | 0.871 |  |

lme per within-network combination: cosine distance = STc vs. STi + confounds.
Task-activity remained in data, schaefer200 parcellation

**Table S30.**
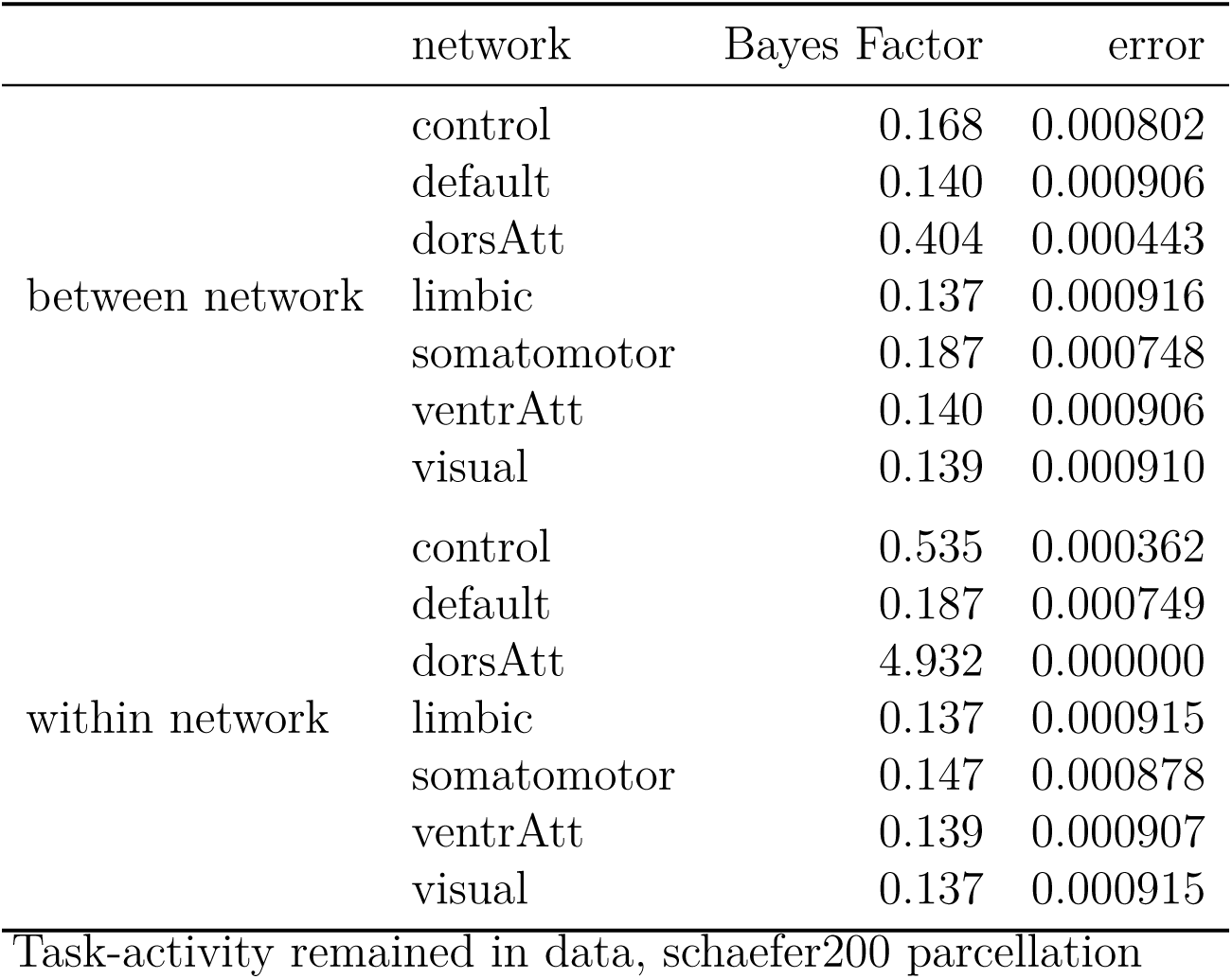
Bayes paired t-test between similarity (cosine distance) of the single task modalitiy pairings.

**Table S31.**
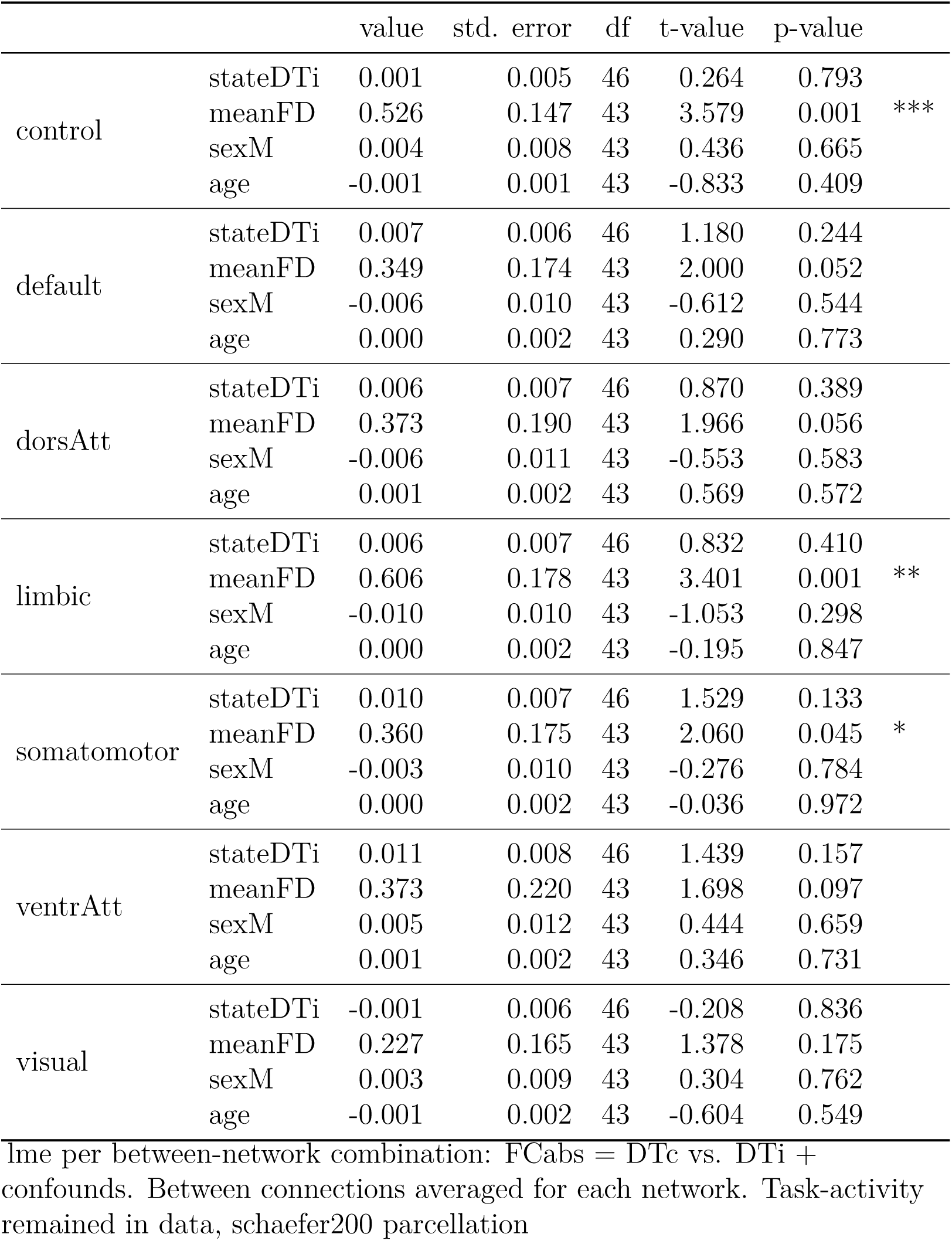
Functional connectivity (absolute) between dual tasks per between-network combination, full model results.

**Table S32.**
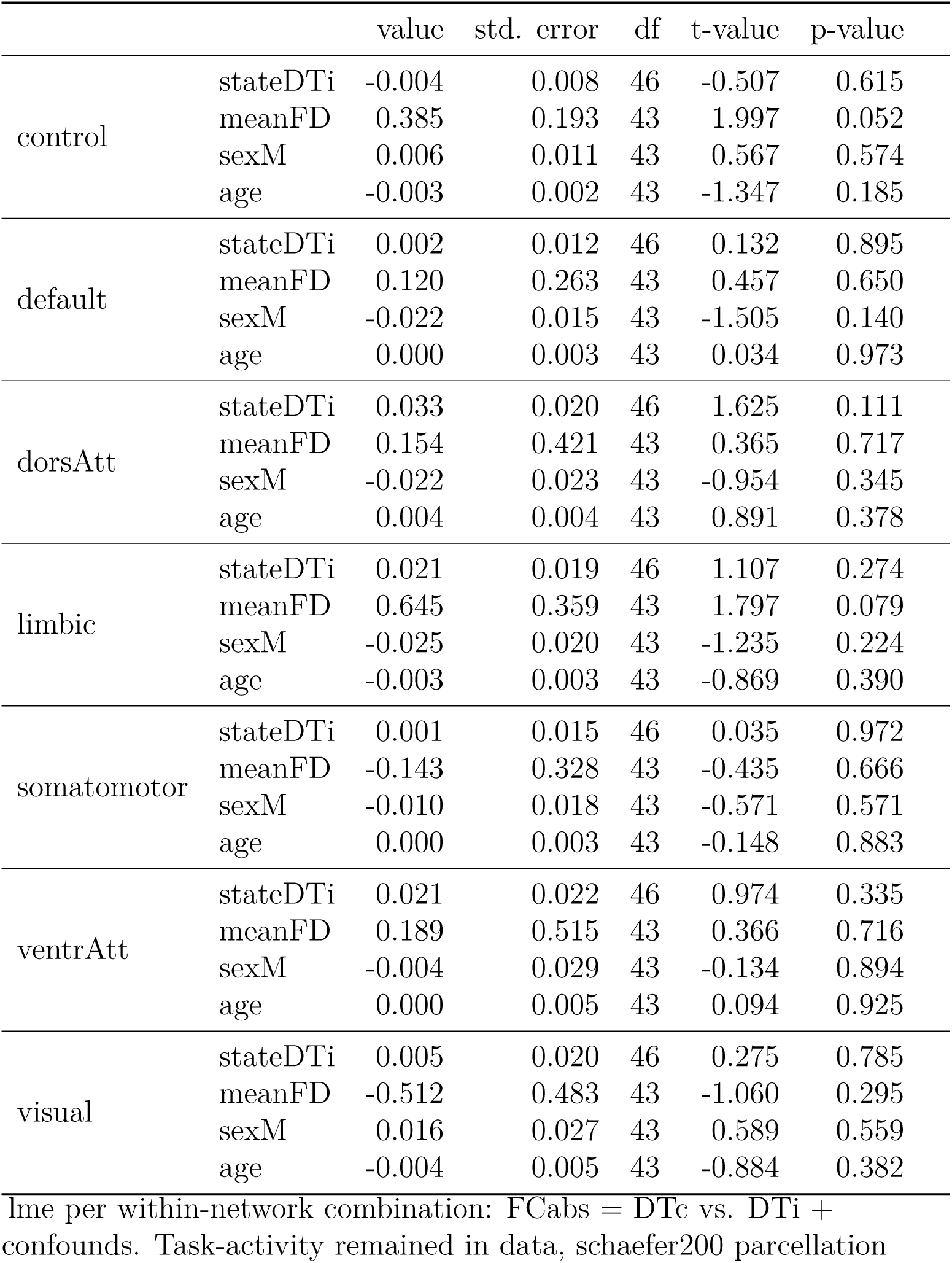
Functional connectivity (absolute) between dual tasks per within-network combination, full model results.

**Table S33.** Bayes paired t-test between absolute functional connectivity of dual task modalitiy pairings.

|  | network | Bayes Factor | error |
| --- | --- | --- | --- |
| between network | control | 0.164 | 0.000628 |
|  | default | 0.310 | 0.000458 |
|  | dorsAtt | 0.229 | 0.000535 |
|  | limbic | 0.222 | 0.000543 |
|  | somatomotor | 0.484 | 0.000358 |
|  | ventrAtt | 0.428 | 0.000384 |
|  | visual | 0.162 | 0.000632 |
| within network | control | 0.180 | 0.000602 |
|  | default | 0.160 | 0.000635 |
|  | dorsAtt | 0.558 | 0.000330 |
|  | limbic | 0.271 | 0.000491 |
|  | somatomotor | 0.158 | 0.000638 |
|  | ventrAtt | 0.251 | 0.000511 |
|  | visual | 0.164 | 0.000627 |
| Task-activity remained in data, schaefer200 parcellation |  |  |  |

### Graph parameters

A common approach in network analysis are the use of graph parameters to describe the network based on graph theory. We calculated two measures, the global efficiency, which quantifies the efficiency of communication in a network as the average of inverse shortest path length (Rubinov & Sporns, 2010) and the network modularity, which describes how well the network can be subdivided into non-overlapping parts. Scripts to calculate both parameters can be found on OSF https://osf.io/w9hsu/.

#### Global modularity

We applied the same algorithm as Hilger and Fiebach (2019) which is based on the Louvain algorithm (Blondel et al., 2008). We used a fixed gamma of 1 and repeated the Louvain community detection 100 times to maximize the modularity parameter *Q*. We selected the ‘negative_asym’ option to again account for negative connections in the matrices, which is suggested by Rubinov and Sporns (2010) in the Brain Connectivity toolbox. As for the global efficiency measure, we calculated the modularity per participant for single, and dual task and for each modality pairing (total of four matrices). Input was the fishers z-transformed functional connectivity matrices. The same mixed model and Bayes paired t-test was used as for the global efficiency.

Similar to the other results, we did not find any evidence for a different modularity between modality pairings (see Figure S3), neither in the mixed model (Table S34) nor in the Baysian paired t-test (Table S35). In sum, we did not find any evidence that the neural network is differently structured in communities for the modality-compatible pairing as for the modality-incompatible pairing.

**Table S34.** Global network modularity (gamma =1)

|  | value | std. error | df | t-value | p-value |  |
| --- | --- | --- | --- | --- | --- | --- |
| mappingincomp | 0.000 | 0.007 | 109 | -0.049 | 0.961 |  |
| taskST | -0.026 | 0.015 | 42 | -1.719 | 0.093 |  |
| sexM | -0.015 | 0.008 | 63 | -1.990 | 0.051 |  |
| run | 0.009 | 0.003 | 42 | 3.509 | 0.001 | ** |
| age | -0.001 | 0.001 | 63 | -1.049 | 0.298 |  |
| meanFD | -0.113 | 0.131 | 42 | -0.857 | 0.396 |  |
| mappingincomp:taskST | 0.001 | 0.009 | 109 | 0.079 | 0.937 |  |

lme model: modularity Q = ST vs. DT \* mapping + confounds.
Task-activity regressed out, schaefer100 parcellation.

**Table S35.** Bayes paired t-test between modalitiy pairings of global modularity score for each combination of task (single and dual task) with gamma = 1.

|  | Bayes Factor | error |
| --- | --- | --- |
| DT | 0.158 | 0.000638 |
| ST | 0.138 | 0.000913 |

Task-activity regressed out, schaefer100 parcellation

**Figure S3.**
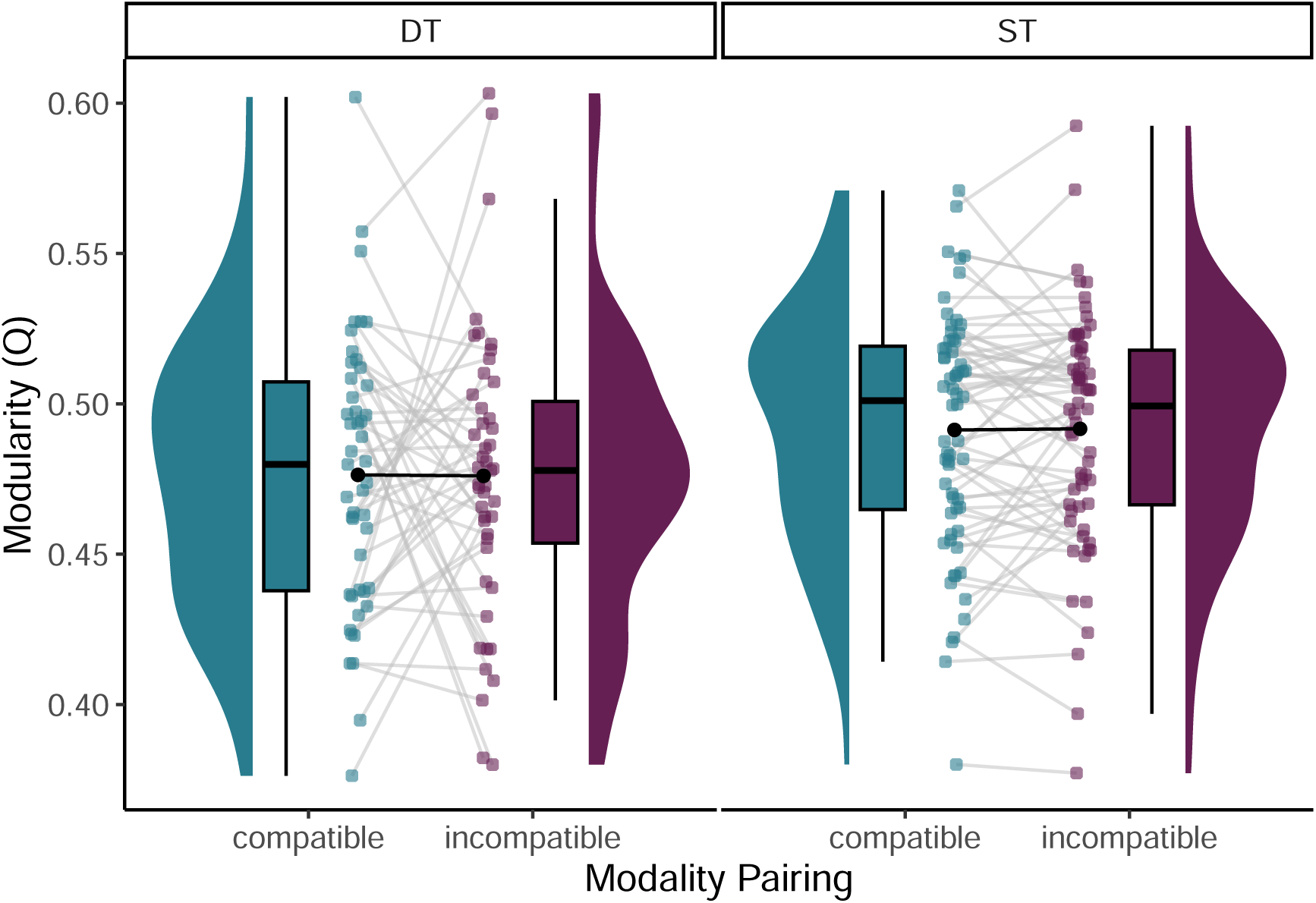
Global modularity per task type comparing modality pairings. Graph shows distribution, boxplot, indidvidual data (color points) and mean value per modality pairing (black). DT = dual task, ST = single task, gamma = 1.

#### Global efficiency

We used the function *efficiency_wei* from the Brain connectivity Toolbox (Rubinov & Sporns, 2010) and applied three different forms of handling negative connections in the FC matrices: filter, absolute and rescale. In the filter option we set the negative connections to NaN, in the absolute option we take only absolute connections and the last option is to rescale all value to the interval [0,1]. As there is no agreement of the best practice how to include negative connections, we applied all three options and compared the results. Input to all three options are the task-related 200x200 fisher’s z-transformed FC matrices per participant, separately for single, dual task and for both modality pairings (total of four matrices). Similar to the other analysis, we were interested in the effect of modality pairing, testing it with a linear-mixed effect model, while controlling for head movement, age, gender and number of runs (formula per analysis option: *globalEfficiency* ∼ *task* ∗ *mapping* + *meanF D* + *run* + *age* + *sex*) and calculated the Bayes Factor for a paired t-test between modality pairing for single and dual tasks.

Figure S4 and the model results in Table S36 showed a significant main effect of task, with higher global efficiency for the dual task, compared to the single task, consistent over the three options. But only for the filter option we see a significant interaction effect between task and modality pairing, which cannot be confirmed by the Bayes Factor being all below 0.6 (compare Table S37). In sum, we did not find evidence for differences in how efficient the network communicate between the modality-compatible and modality-incompatible pairing, neither during single, nor during dual task.

**Table S36.** Global network efficiency per negative-weight treatment.

|  |  | value | std. error | df | t-value | p-value |  |
| --- | --- | --- | --- | --- | --- | --- | --- |
| abs | (Intercept) | 0.535 | 0.027 | 109 | 19.591 | 0.000 | *** |
|  | taskST | -0.127 | 0.013 | 42 | -9.853 | 0.000 | *** |
|  | mappingincomp | 0.005 | 0.003 | 109 | 1.446 | 0.151 |  |
|  | meanFD | 0.093 | 0.109 | 42 | 0.856 | 0.397 |  |
|  | run | -0.009 | 0.002 | 42 | -4.124 | 0.000 | *** |
|  | age | -0.001 | 0.001 | 63 | -0.630 | 0.531 |  |
|  | sexM | 0.000 | 0.006 | 63 | 0.043 | 0.965 |  |
|  | taskST:mappingincomp | -0.008 | 0.005 | 109 | -1.679 | 0.096 |  |
| filt | (Intercept) | 0.407 | 0.019 | 109 | 21.807 | 0.000 | *** |
|  | taskST | -0.074 | 0.009 | 42 | -8.384 | 0.000 | *** |
|  | mappingincomp | 0.004 | 0.002 | 109 | 2.182 | 0.031 | * |
|  | meanFD | 0.046 | 0.074 | 42 | 0.618 | 0.540 |  |
|  | run | -0.007 | 0.002 | 42 | -4.902 | 0.000 | *** |
|  | age | 0.000 | 0.001 | 63 | -0.408 | 0.684 |  |
|  | sexM | 0.002 | 0.004 | 63 | 0.588 | 0.559 |  |
|  | taskST:mappingincomp | -0.006 | 0.003 | 109 | -2.293 | 0.024 | * |
| resc | (Intercept) | 0.471 | 0.024 | 109 | 19.297 | 0.000 | *** |
|  | taskST | -0.084 | 0.013 | 42 | -6.586 | 0.000 | *** |
|  | mappingincomp | 0.005 | 0.004 | 109 | 1.060 | 0.292 |  |
|  | meanFD | 0.151 | 0.100 | 42 | 1.511 | 0.138 |  |
|  | run | -0.006 | 0.002 | 42 | -2.586 | 0.013 | * |
|  | age | 0.000 | 0.001 | 63 | -0.424 | 0.673 |  |
|  | sexM | 0.003 | 0.005 | 63 | 0.665 | 0.509 |  |
|  | taskST:mappingincomp | -0.005 | 0.006 | 109 | -0.824 | 0.412 |  |

lme per negative-weight treatment: efficiency = ST vs. DT \* mapping + confounds. Task-activity regressed out, schaefer200 parcellation. abs = absolute values, filt = negative connections are filtered out, resc = rescale connections.

**Table S37.**
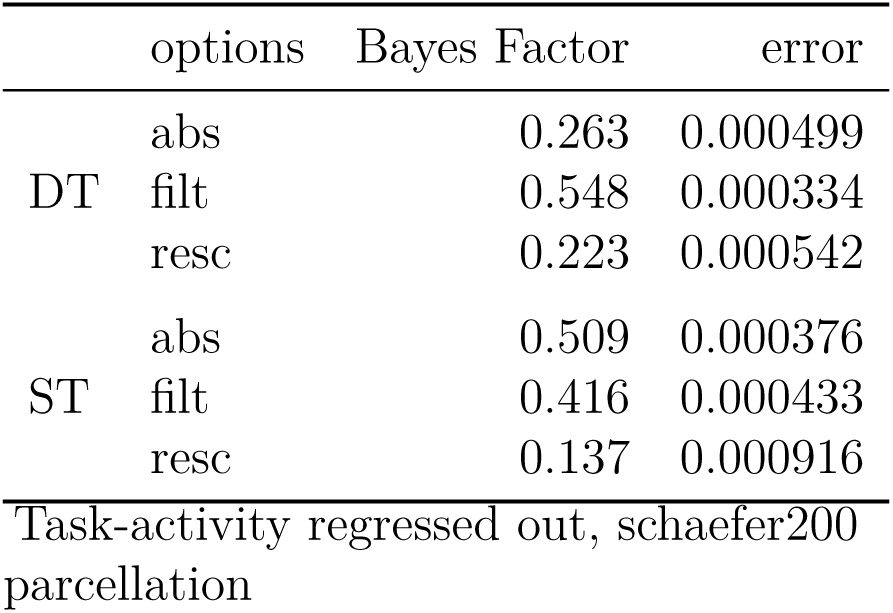
Bayes paired t-test between modalitiy pairings of global efficiency score per negative-weight treatment.

**Figure S4.**
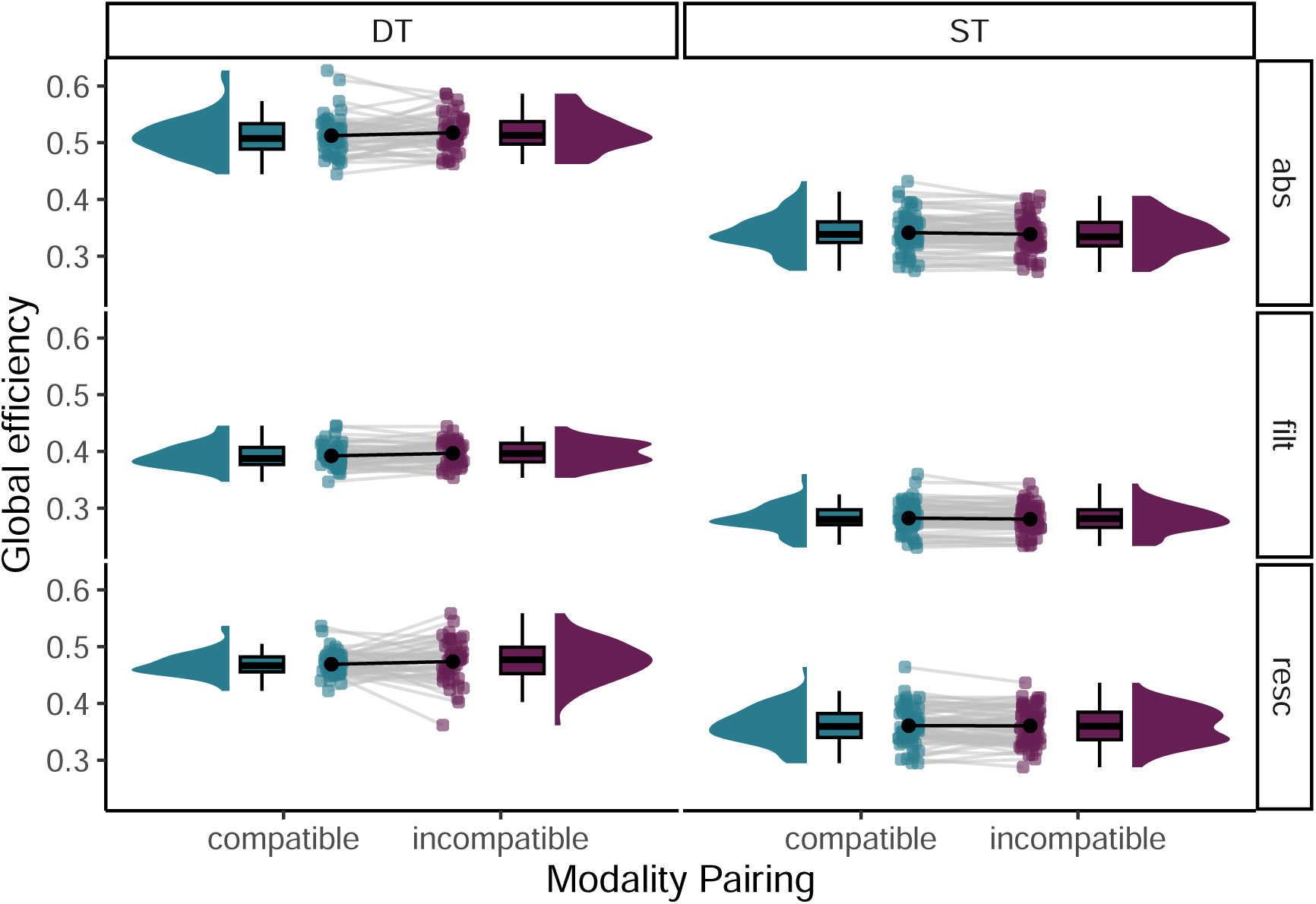
Global efficiency per task type and analysis option comparing modality pairings. Graph shows distribution, boxplot, indidvidual data (color points) and mean value per modality pairing (black). DT = dual task, ST = single task, abs = absolute FC matrix, filt = filtered for only positive weigths, resc = rescale FC matrix to interval [0,1].

